# Biophysically inspired mean-field model of neuronal populations driven by ion exchange mechanisms

**DOI:** 10.1101/2021.10.29.466427

**Authors:** Giovanni Rabuffo, Abhirup Bandyopadhyay, Carmela Calabrese, Kashyap Gudibanda, Damien Depannemaecker, Lavinia Mitiko Takarabe, Sourin Chatterjee, Maria Luisa Saggio, Mathieu Desroches, Anton Ivanov, Marja-Leena Linne, Christophe Bernard, Spase Petkoski, Viktor K. Jirsa

## Abstract

Whole-brain simulations are a valuable tool for gaining insight into the multiscale processes that regulate brain activity. Due to the complexity of the brain, it is impractical to include all microscopic details in a simulation. Hence, researchers often simulate the brain as a network of coupled neural masses, each described by a mean-field model. These models capture the essential features of neuronal populations while approximating most biophysical details. However, it may be important to include certain parameters that significantly impact brain function. The concentration of ions in the extracellular space is one key factor to consider, as its fluctuations can be associated with healthy and pathological brain states. In this paper, we develop a new mean-field model of a population of Hodgkin–Huxley-type neurons, retaining a microscopic perspective on the ion-exchange mechanisms driving neuronal activity. This allows us to maintain biophysical interpretability while bridging the gap between micro- and macro-scale mechanisms. Our model is able to reproduce a wide range of activity patterns, also observed in large neural network simulations. Specifically, slow-changing ion concentrations modulate the fast neuroelectric activity, a feature of our model that we validated through in vitro experiments. By studying how changes in extracellular ionic conditions can affect whole-brain dynamics, this model serves as a foundation to measure biomarkers of pathological activity and provide potential therapeutic targets in cases of brain dysfunctions like epilepsy.

## 1 Introduction

Large-scale brain dynamics can be studied in silico with network models [1]. Local activity can be represented by neural mass models [2], which coupled together through synapses, time delays, and noise [3–6] allow the emergence of whole-brain activity that can be linked to empirical neuroimaging data [7]. In the context of large-scale simulations, these models have been employed for the study of resting-state brain activity (i.e., in the absence of any stimulus or task) in several mammalian species [8–10], for the analysis and identification of chaos in brain signals [11, 12], the study of epileptic seizure genesis and propagation [13–18], among other applications.

At the mesoscopic level, the observable properties of a neuronal ensemble are generally explained by statistical physics formalism of mean-field theory [19–22]. Mean-field models demonstrated a predictive value for studying the mesoscopic dynamics of neuronal populations [23], providing statistical descriptions of neuronal networks [2, 19, 24–29], which can be used to address questions related to network-level mechanisms [12, 24, 30]. In general, neural mass models have a low enough number of parameters to be tractable and provide general intuitions regarding mechanisms underlying complex neuronal activity [31–36]. For example, statistical population measures, such as the firing rate, can be used to assess mesoscopic dynamics [1, 7, 31, 36–41].

Recently, a class of these models, called next-generation neural mass models [42], has been developed based on an analytical approach introduced by [25] that allowed for the exact derivation of mean field parameters for a population of quadratic integrate-and-fire (QIF) neurons. These can be linked to EEG/MEG oscillations [43], including epipeltic seizures [44], and have been used to study various aspects of the whole-brain dynamics such as the low-dimensional manifold of the resting state [45, 46], aging [47] and neural signatures of consciousness [48]. Number of works dealt with the introduction of biologically realistic aspects in the mostly phenomenological neural mass model derived in [25]. These included short-term synaptic plasticity [49–51], spike frequency adaptation [52, 53], spike timing-dependent plasticity [54], synaptic delay [29], random connectivity and noise [55–57], as well as an extension of the conductance-based neurons with a recovery variable [58–60].

Although it is practically inconvenient to reproduce the entire complexity of a neural mass, including all known biophysical parameters, it may be important to include parameters that can have widespread and general effects on neuronal activity [34]. The concentrations of *Na*^+^, *K*^+^, *Ca*^2+^, and *Cl*^*−*^ ions in the extracellular space are key parameters to consider. Extracellular ion concentrations change dynamically in vivo as a function of the brain state, for example, between arousal and sleep [61, 62]. Changes in extracellular ion concentrations can induce fluctuations in the spontaneous activity [63], in particular in the awake state [64, 65] as well as to switch one brain state to another [61]. In particular, the extracellular potassium concentration [*K*^+^]_*ext*_ plays a central role. Transient changes in [*K*^+^]_*ext*_ can have large effects on cell excitability and spontaneous neuronal activity [66–68], a result consistently reported in modeling studies [69–71]. Increases in [*K*^+^]_*ext*_ are tightly controlled by astrocytes [72], which can efficiently pump out [*K*^+^]_*ext*_ and distribute it via their syncytium to prevent hyperexcitability [73–77]. The saturation or lack of efficiency of these buffering mechanisms is often linked to a pathological state. Detailed single neuron models demonstrate that continuous increases in [*K*^+^]_*ext*_ can lead to different firing states, from tonic to bursting, to seizure-like events and depolarization block [68, 78]. In such detailed models, the buffering action of astrocytes is represented by a parameter named [*K*^+^]_*bath*_ [68, 73, 74, 78, 79]. This parameter may also account for the potassium concentration in the perfusion bath registered in vitro. Our goal is to extend the use of ion-concentration variables at the neural mass level, to be incorporated into whole-brain modeling.

In this study, we applied a mathematical formalism to estimate the mean-field behavior of a large neuronal ensemble, taking into account the ion exchange between the intracellular and extracellular space. Large networks of biophysically realistic neurons display complex behavior and–likely–do not satisfy the typical conditions required for deriving the mean-field dynamics (e.g., the Lorentzian Ansatz [80]). In this work, relying on approximations and heuristic arguments, we derive a new neural mass model of a large population of an all-to-all coupled network of heterogeneous Hodgkin-Huxley-like neurons operating near synchrony (Fig 1). While the derivation is not exact, our model captures the mean-field behavior of connected neuronal populations operating in a synchronous regime, also displaying emergent dynamics not present in the single-neuron mode, as we show comparing our neural mass model outcome with the simulated activity from a large number of connected neurons. Our model faithfully characterizes the slow modulation of local field potential fluctuations depending on ion concentrations, which we confirm through in vitro experiments. Considering different parameter configurations, we identify the mesoscopic states of the connected neuronal population, in various dynamical regimes that can be linked to different healthy and pathological states, and be of use in large-scale brain simulations. This approach establishes a link between the biophysical description at the cellular scale and the dynamics observable at the mesoscopic scale, enabling the study of the influence of changes in extracellular ionic conditions on whole brain dynamics in health and disease.

**Fig. 1.**
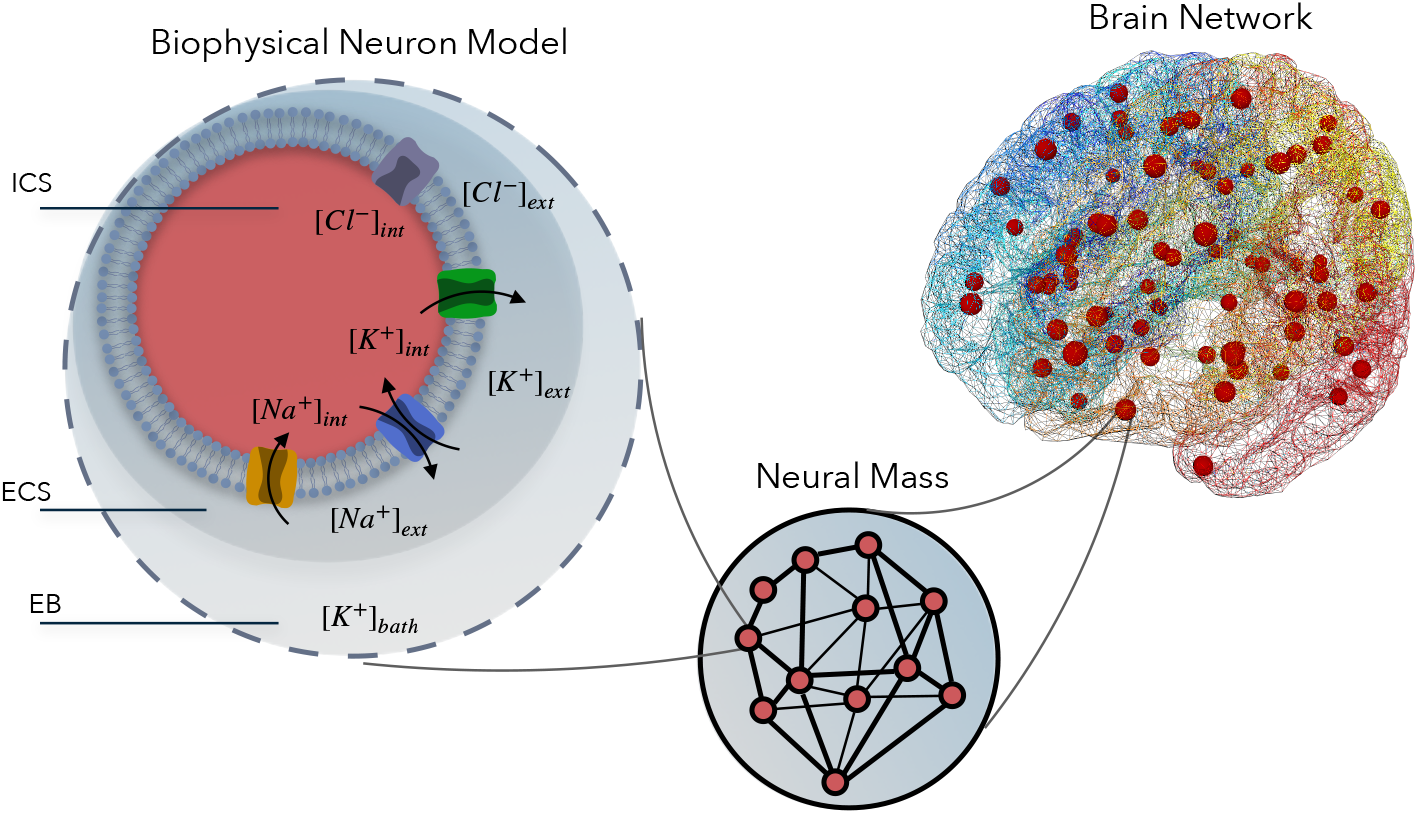
Biophysically inspired neural mass model: Schematic diagram of the ion channel mechanism in extracellular and intracellular space in the brain. A biophysical model of a single neuron consists of three compartments (left panel): the intracellular space (ICS; in red), the extracellular space (ECS; in dark gray), and the external bath (EB; in light gray). The ion exchange across the cellular space occurs through the ion channels: *Na*^+^ gets inside the ICS (yellow channel), *K*^+^ gets out (green channel), the flow of *Cl*^*−*^ can be bidirectional (purple channel); for the pump (blue), *Na*^+^ gets out and *K*^+^ gets into the ICS. A population of interacting neurons sharing the same [*K*^+^]_*bath*_ concentration forms a local neural mass (middle panel), for which we model the mean-field equations in this work. Brain network model (right panel) with the activity of each brain region represented by neural masses.

## Results

### 1.1 Biophysically inspired mean-field model

In this work, we derived a mean-field model describing the activity of a neural mass regulated by ion exchange mechanisms at the cellular level (Fig. 1). This model establishes a computationally accessible baseline that allows large-scale brain activity to be understood in terms of a few key biophysical details regulating the micro-scale mechanisms. The mean-field model equations (38) were derived by approximating a locally homogeneous network of Hodgkin–Huxley (HH) type neurons operating near synchrony, in the thermodynamic limit of an infinitely large population (see Methods section). By locally homogeneous, we mean that all neurons in the population are assumed to share the same extracellular and intracellular ionic environment and are connected with identical coupling rules, allowing us to treat the population as uniform with respect to ion dynamics and connectivity.

The single neuron equations (1) were previously derived in [78] as a simplified version of HH neurons which includes three compartments: an intracellular space (ICS), and extracellular space (ECS) and an external bath (EB) in communication through ion-fluxes (Fig. 1, left). The decoupled single-neuron equations exhibit a range of activity patterns including bursting behavior, tonic spiking, seizure-like events, sustained ictal activity, and depolarization block (Fig. 2.a). As we will describe in more detail in the next sections, also the mean-field model exhibits these activity patterns, together with new complex behaviors emergent from the network interactions at the neural mass level. The variables describing a single neuron are characterized by a fast compartment, including membrane potential and gating variable fluctuations, and a slow compartment, which includes the slow-fluctuating ion concentrations (Fig. 2.b). The mean-field model inherits the single neuron biophysical parameters (Table1), which can be tuned to match different neuron types. For example, excitatory or inhibitory neurons can be characterized by tuning the reversal potential to high (e.g., *E*_*syn*_ = 0mV) and low (e.g., *E*_*syn*_ = −80mV) values, respectively. It was previously established that a system of all-to-all coupled neuronal equations can be solved exactly in the thermodynamic limit (i.e., infinite neurons limit) if the single neuron membrane potential equation is a quadratic function and if the instantaneous distribution of membrane potentials of neurons in a population is described by a Lorentzian [25]. In our case, for any fixed value of the gating and potassium variables, the membrane potential equation resembles a cubic function (Fig. 2.c).

**Table 1.** List of parameters and their values used for the single-neuron simulation.

| Parameter | Symbol | Value |
| --- | --- | --- |
| Membrane capacitance | $C_m$ | 1nF |
| Gating time constant | $\tau_n$ | 4ms |
| Chloride conductance | $g_{Cl}$ | 7.5nS |
| Maximal potassium conductance | $g_K$ | 22nS |
| Maximal sodium conductance | $g_{Na}$ | 40nS |
| Potassium leak conductance | $g_{K,l}$ | 0.12nS |
| Sodium leak conductance | $g_{Na,l}$ | 0.02 nS |
| Intracellular volume | $\omega_i$ | 2160$\mu\text{m}^3$ |
| Extracellular volume | $\omega_{ext}$ | 720$\mu\text{m}^3$ |
| Intra/extra-cellular volume ratio | $\beta = \omega_i/\omega_{ext}$ | 3 |
| Conversion factor | $\gamma$ | 0.04 mol/C |
| Diffusion rate | $\epsilon$ | 0.001mHz |
| Maximal Na/K pump current | $\rho$ | 250pA |
| Initial concentration of Extracellular K | $[K^+]_{0,ext}$ | 4.8 mmol/m<sup>3</sup> |
| Initial concentration of Intracellular K | $[K^+]_{0,int}$ | 130 mmol/m<sup>3</sup> |
| Initial concentration of Extracellular Na | $[Na^+]_{0,ext}$ | 138 mmol/m<sup>3</sup> |
| Initial concentration of Intracellular Na | $[Na^+]_{0,int}$ | 16 mmol/m<sup>3</sup> |
| Initial concentration of Extracellular Cl | $[Cl^-]_{0,ext}$ | 112 mmol/m<sup>3</sup> |
| Initial concentration of Intracellular Cl | $[Cl^-]_{0,int}$ | 5 mmol/m<sup>3</sup> |

**Fig. 2.**
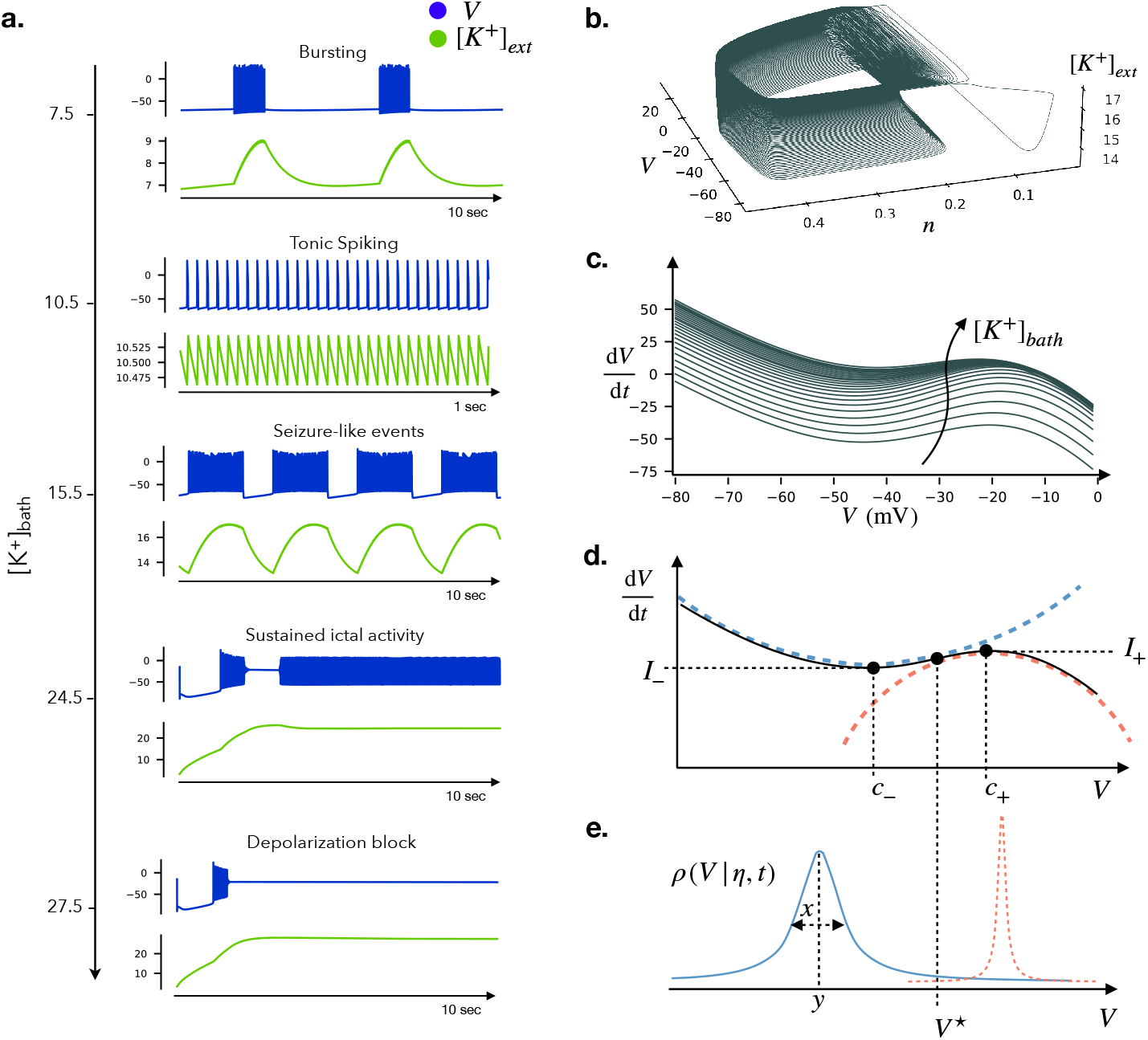
Single Hodgkin-Huxley type neuron model: **(a)** Different patterns of electrophysiological activities previously identified in [78] are also reproduced in our parameter setting by varying the potassium concentration in the external bath [*K*^+^]_*bath*_. The membrane potential *V* is measured in mV and the ion concentrations in *m*mol/m^3^. **(b)** Phase space trajectory of the seizure-like event simulation ([*K*^+^]_*bath*_ = 15.5). Fast oscillations occur in a fast sub-system identified by the membrane potential *V* and the gating variable *n*. The oscillations of the slow subsystem, here captured by the potassium concentration in the extracellular space [*K*^+^]_*ext*_, enable the transition to bursting. **(c)** Fixing the value of the state variables *n*, Δ[*K*^+^]_*int*_ and [*K*^+^]_*g*_ as constants, the membrane potential equation resembles a cubic function for different values of [*K*^+^]_*bath*_. We can model this function as a step-wise quadratic approximation, corresponding to two parabolas with vertices at coordinates (*c*_*−*_, *I*_*−*_) and (*c*_+_, *I*_+_) and curvature *R*_*−*_ and *R*_+_ respectively **(d).** The two parabolas meet at an intersection point *V* ^⋆^ where the membrane potential equation changes curvature. **(e)** At each time, we assume that the membrane potential of a neuronal population is distributed according to a Lorentzian centered at *y* = *y*(*η, t*) and with width *x* = *x*(*η, t*), for each value of the excitability *η* (Lorentzian Ansatz). In the case depicted, the cubic function meets the zero for *V* < *V* ^⋆^, the neuronal population is described by the Lorentzian distribution in blue in the steady-state solution, and the neuronal dynamics is governed by the positive parabola according to the continuity equation. In the case where the derivative of the membrane potential crosses zero for *V* > *V* ^⋆^ (e.g., if the cubic function is shifted up by adding a constant current to the membrane potential derivative), the population is described by the red distribution in the steady state, and the continuity equation is governed by the negative parabola equation. Cases where the cubic function meets the zero in more than one point are not well described by this approximation (see Section *Steady-state solution and Lorentzian Ansatz*

Therefore, to proceed with the mean-field reduction, we make several key assumptions: 1) the cubic-like profile can be described by a step-wise quadratic function corresponding to two parabolas with opposite curvature (Fig. 2.d); 2) The membrane potential distribution of HH type neurons is described at each time by a Lorentzian (Fig. 2.e), which is a good approximation in situations where neurons are operating close to synchrony (e.g., Fig. 3.a), but is not a suitable description for other dynamical states better described by a bimodal distribution (e.g., Suppl. Fig. S1.a, orange distribution); 3) We assume that the potassium concentrations, both intracellular (Δ[*K*^+^]_int_) and extracellular (through the buffering variable [*K*^+^]_*g*_), are homogeneous across the neuronal population. This is justified physiologically by the rapid redistribution of ions through diffusion and electrochemical gradients, which enforce near-instantaneous equilibration at the mesoscopic scale. As such, assigning separate compartments to each neuron is neither practical nor biologically meaningful in this context. We assume that the potassium concentrations, both intracellular (Δ[*K*^+^]_int_) and extracellular (through the buffering variable [*K*^+^]_*g*_), are homogeneous across the neuronal population. This is justified physiologically by the rapid redistribution of ions through diffusion and electrochemical gradients, which enforce near-instantaneous equilibration at the mesoscopic scale. As such, assigning separate compartments to each neuron is neither practical nor biologically meaningful in this context; 4) We assume that the gating variable *n*, which governs potassium conductance, can be treated as a population-averaged variable. This allows us to describe the neuronal ensemble using a reduced set of collective (mean-field) variables.

**Fig. 3.**
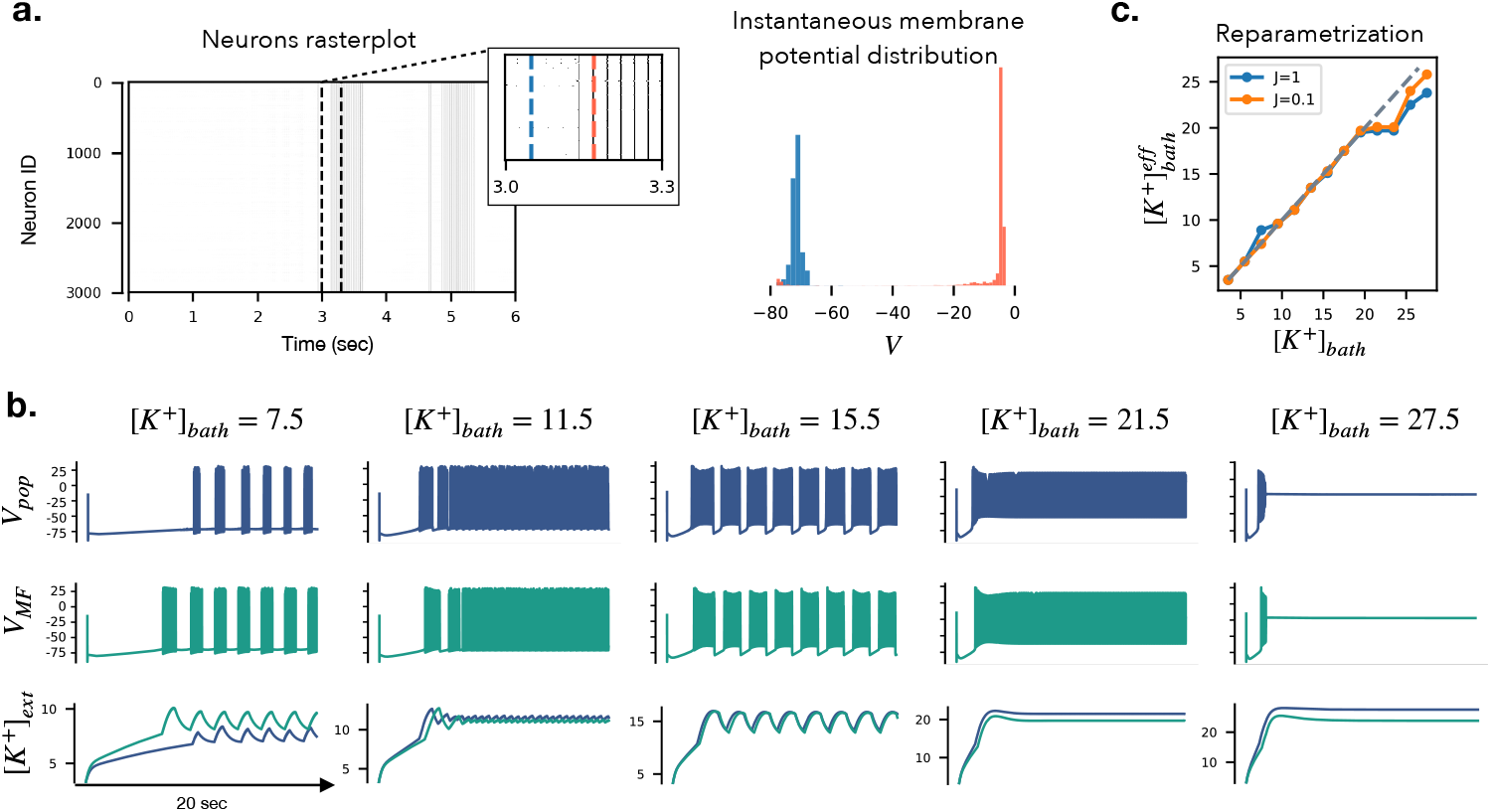
Mean-field model versus Neuronal network across dynamical regimes: Example raster plot of a population of *N* = 3000 all-to-all coupled HH-type neurons displaying sub- and supra-threshold dynamics that can be well described by a Lorentzian distribution. **(b)** For several [*K*^+^]_*bath*_ values we simulated the activity of a population of *N* = 3000 all-to-all coupled HH-type neurons across dynamical regimes (parameters in Table S2). We compare the mean membrane potential *V*_*pop*_ and external potassium [*K*^+^]_*ext*_ of such population (in blue) with the results obtained using the mean-field model equations (in green). To properly match the slow timescale of the population, we defined an effective value of the potassium concentration in the bath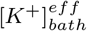, represented in panel **(c)** (the dashed line represents the identity).

Using these key approximations, we derived the primary result of this work: a closed form for the neural mass equation (Eq.38) expressed in terms of mean-field variables.

### 1.2 Comparison with neural network simulations

The accuracy of the mean-field approximation is first validated by comparing it to the simulation of a large network of coupled HH-type neurons. The simulation of Hodgkin–Huxley type single neuron dynamics (Eq. (1)) driven by an ion-exchange mechanism, has revealed that the parameter [*K*^+^]_*bath*_ has a significant impact on the dynamics (Fig. 2.a). Thus, for different [*K*^+^]_*bath*_ we simulated a network of *N* = 3000 coupled neurons described by Eq. (1), coupled as inEq.(9) with synaptic strength *J* = 1 and heterogeneous excitability distributed with mean 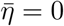 and width Δ = 1.

Using the mean-field equations Eq.(38), we show that the mean-field model membrane potential *V*_*MF*_ matches qualitatively the average membrane potential *V*_*pop*_ of the neuronal population (Fig. 3.b). We stress that *V*_*MF*_ was obtained by solving 5 coupled equations Eq.(38), while *V*_*pop*_ was obtained by averaging the solution of 4 × 3000 equations (1) coupled as in Eq.(9). This result shows that the mean-field model can be used to simulate several regimes of activity and emulate the average dynamics of a large neuronal population in regimes where the membrane potentials’ distribution is unimodal and can be reasonably approximated by a Lorentzian. These regimes include activities such as spike trains, tonic spiking, bursting, seizure-like events, status epilepticus-like events, and depolarization block (such as in Fig. 3.b). Notice that in such regimes the population average displays dynamics qualitatively similar to the single neuron simulation (Fig. 2.b). However, due to the network effects in the population, there is a frequency shift compared to the single neuron dynamics (for example, the bursts in the [*K*^+^]_*bath*_ = 7.5 regime are ∼3 times faster in the population Fig. 3.b than in the single neuron Fig. 2.a).

A frequency shift is also present in the mean-field model compared to the population dynamics. However, the definition of an effective 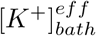 can adjust this frequency shift (Fig. 3.c). In Fig. 3.c we report the 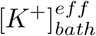 values obtained via visual inspection of the match with the population dynamics for two values of synaptic strength *J* = 0.1 and *J* = 1, and used in (Fig. 3.b). Also note that the gating variable *n* is treated as microscopic in the neural network, while in the derivations for the mean-field it is considered as a mesoscopic and identical for the whole population. This is likely responsible for some of the discrepancies between the two modalities. [49–51]

### 1.3 Comparison with in vitro experiments

We rearranged Figure 4 to better highlight the range of dynamics involving extracellular potassium concentration and bursting activity. The neural mass model (Fig. 4.a) reproduces a periodic behavior of the extracellular potassium concentration, with fast voltage bursts riding on top (we used the parameters *C*_*m*_ = 16, *τ*_*n*_ = 8, *ϵ* = 0.0001, *γ* = 0.00025). A similar pattern is observed in the in vitro recordings (Fig. 4.c), although we emphasize that these are AC-coupled LFP traces. As such, slower components of the membrane potential — including jumps during bursting — are filtered out and not visible in the LFP recordings. We do not attempt to classify the bursting pattern in either data or simulations. To compare with experimental timescales, we also simulate a network of *N* = 3000 HH-type neurons (Fig. 4.b). The parameters were adjusted for computational efficiency, yielding a shorter bursting period, but preserving the key qualitative feature: modulation of fast activity by slow potassium dynamics. More complex patterns can emerge in vitro as well. For example, in Fig. 4.d, we observe a progressive slowing of the burst frequency, which our deterministic model does not capture. We hypothesize that such behavior may arise from slow parameter drifts or noise-driven transitions between metastable regimes — effects that are not included in the present model, but could be explored by introducing noise or varying parameters such as [*K*^+^]_bath_ or the nullcline geometry (see e.g., [81]). Finally, Fig. 4.e shows an emergent regime from the mean-field model with isolated bursts in the up state, qualitatively resembling some features of the in vitro activity.

**Fig. 4.**
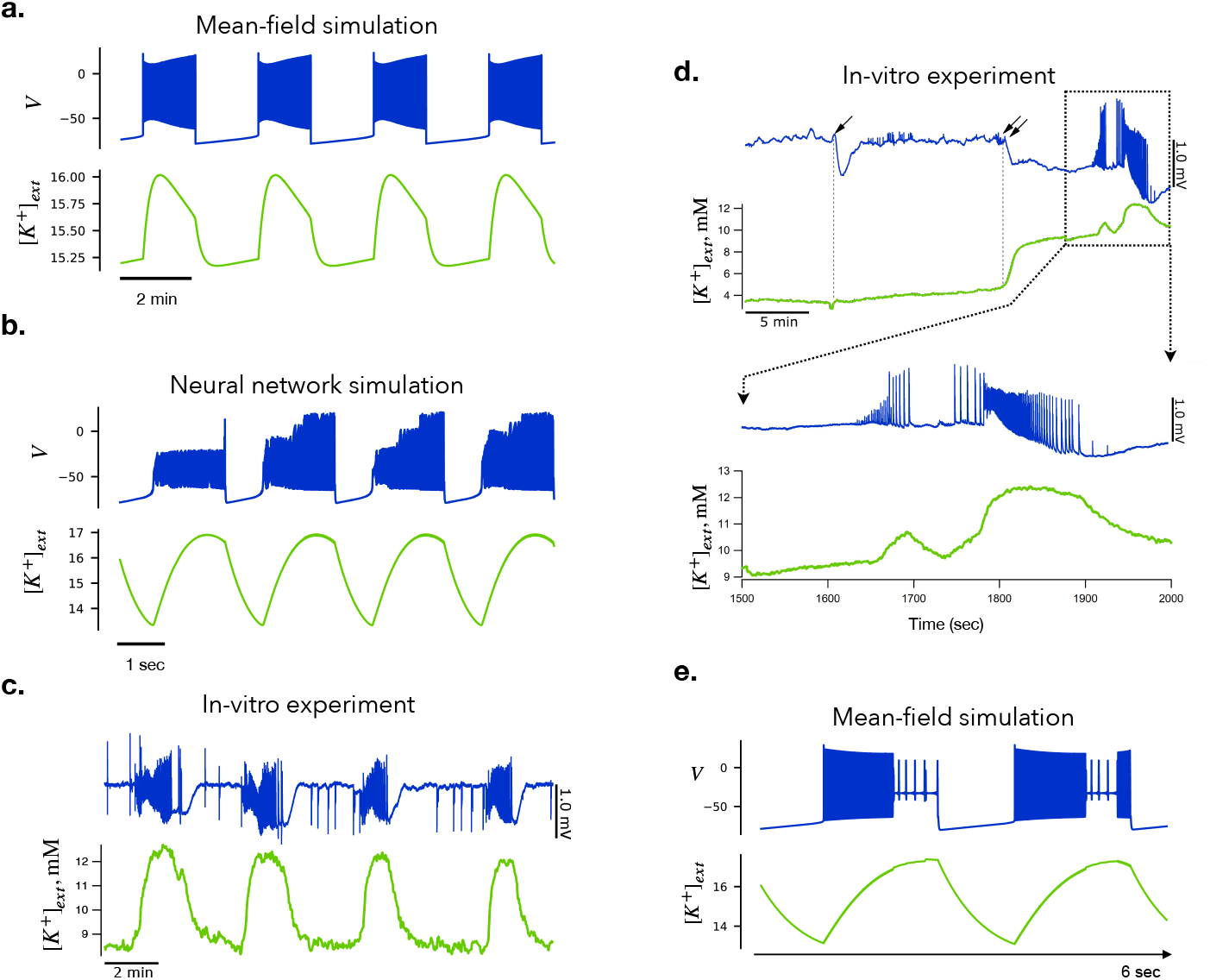
Comparison of numerical results and in vitro experiments: **(a)** Neural mass model showing slow periodic fluctuations in extracellular potassium concentration (green) with voltage bursts (blue) riding on top. **(b)** Network simulation of *N* = 3000 coupled HH neurons exhibiting a similar bursting pattern at a shorter timescale (due to rescaled parameters). **(c)** In vitro recording showing LFP (blue, AC-coupled) and extracellular potassium concentration (green), exhibiting slow periodic fluctuations and bursting activity. **(d)** In vitro trace showing more complex dynamics — after a shift to a high-potassium state (marked by arrows), the burst frequency slows down progressively. **(e)** Simulation from the mean-field model showing an emergent bursting regime with isolated events in the up state, not seen at the single-neuron level.

### 1.4 Bifurcation analysis: emergent network states and multistability

A previous study has established that varying the external potassium concentration ([*K*^+^]_*bath*_) induces significant changes in the dynamical behavior of the single neuron model [78]. Within a specific range of [*K*^+^]_*bath*_, the system exhibits multistability, a phenomenon crucially analyzed using bifurcation analysis. This approach allows us to assess system stability and qualitative behavior independently of initial conditions.

We here applied multiple-timescale bifurcation analysis to the neural mass model (38). Indeed, we treated the variables Δ[*K*^+^]_*int*_ and [*K*^+^]_*g*_ as slow compared to the other variables (due to the relative magnitude of parameters *ε, γ* and *ω*_i_) and we considered the so-called fast subsystem. This corresponds to the limiting system in which slow variables’ dynamics are frozen and slow variables become parameters that are varied to discover the corresponding bifurcation structure (see Fig. 5.a). This analysis unveiled complex regimes within the neural mass model, including distinct regions of multistability. The oscillatory behavior observed in the fast subsystem occurs within the range bounded by the Saddle-Homoclinic (SH), the Hopf2, and the Fold of Limit Cycles FLC1 curves (grey area).

**Fig. 5.**
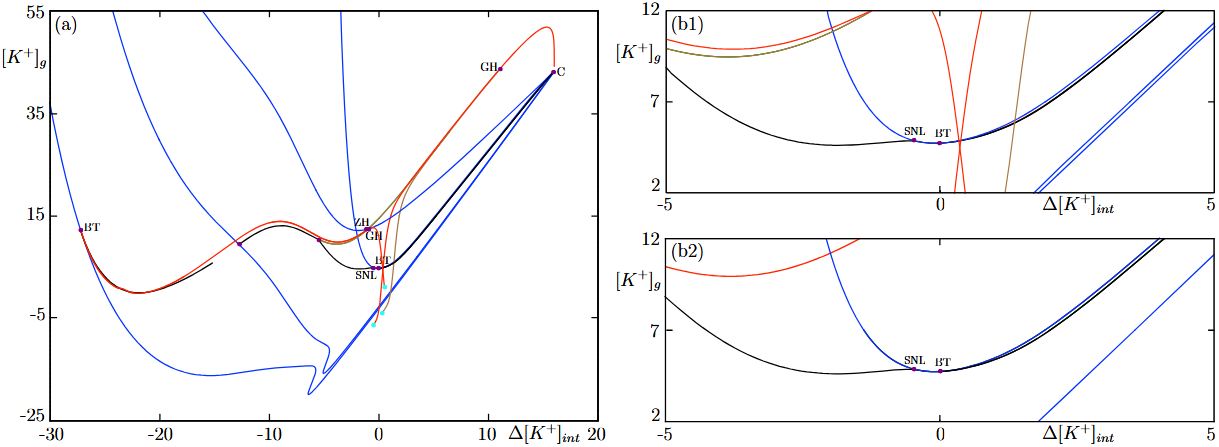
Fast Subsystem Bifurcation Diagram in Two Parameters: **(a)** The bifurcation diagram shows the behavior of the fast subsystem of system (38) to the slow variables Δ[*K*^+^]_*int*_ and [*K*^+^]_*g*_, which are parameters in the fast-subsystem limit. Each curve represents parameter values for which specific bifurcations occur, obtained through a numerical continuation algorithm. Saddle Node (SN) bifurcations are shown in blue, Saddle Homoclinic (SH) bifurcations in black, Fold Limit Cycle (FLC) bifurcations in green, and Hopf bifurcations in red. Colored dots indicate codimension-2 points. In particular, a Bogdanov-Takens (BT) point and a SNL point also identified in the single neuron model, and epileptor model. Note that this diagram is quite involved and we do not claim that the present version is complete, however it is representative of the complexity of the bifurcation structure of the mean-field model. (b1-b2) Comparison of the fast subsystem bifurcation diagrams for the single neuron model and the neural mass model. The left side shows a zoomed-in version of the diagram from panel (a) to facilitate comparison with the single neuron model diagram on the right. Both diagrams use the same color coding as described above. This comparison highlights the similarities and differences in the bifurcation structures of these two models and indicates emergent structures due to interactions within the network.

The qualitative dynamics, illustrated in Fig. 3.b, are influenced by temporal variations in the slow variables and the specific bifurcations encountered in the fast subsystem diagram. They occur in the surroundings of the Saddle-Node-Loop (SNL) point (marked with a yellow star in Fig. 5.b), which is the region of local topological equivalence between the single neuron and the neural mass model. For instance, in our previous study on single neuron dynamics [78], we have shown that seizure-like events arise from transitions between fixed point and limit cycle dynamics and vice versa through SN and SH bifurcations (REF). Similarly, in the neural mass model, seizure-like events (Fig. 3.b at [*K*^+^]_*bath*_ = 15.5) also result from transitions involving these bifurcations. Throughout the burst phase, the slow variables remain constrained between the SN2 and SH curves.

However, further away from the SNL point, the topological equivalence between single neuron and neural mass models is broken. In summary, our neural mass model displays a richer bifurcation structure than the single neuron one (Fig. 5.b, right versus left panels), with new complex activity regimes, which is also supported by the simulation of new behaviors as in Fig. 4.e.

While these results confirm that the neural mass model is not simply mirroring the single neuron one, a detailed numerical exploration of the activity regimes for a population of all-to-all coupled HH-like neurons is required to confirm the accuracy of the neural mass model in representing population behavior emerging from network interactions.

### 1.5 Coupled Neural Masses

Finally, we show that the presented mean-field model can be used to perform large-scale network simulations of brain activity e.g., to be used in the context of brain stimulation or epilepsy. To do so, we coupled six neural masses via long-range structural connections with random weights (Fig. 6.a). Each population (network node) is described by the mean-field model derived in this article (Eq. (38)). Nodes *A, B, C, E, F* are tuned to a healthy regime with [*K*^+^]_*bath*_ = 5.5 which do not show pathological bursts when decoupled from other external inputs. Node *D* is tuned to a pathological regime, with [*K*^+^]_*bath*_ = 15.5. We run a simulation of the system by setting the global coupling parameter to *G* = 0 (Fig. 6.b) and *G* = 100 (Fig. 6.c). In the first case, the system is effectively decoupled and, as expected, the only node displaying pathological activity is node *D*. However, in the second case, when *G* is increased, the system is coupled and the pathological fluctuations of node *D* differentially affect the activity of all other nodes at the whole-network level. The evoked activity in downstream regions is induced by a network reorganization phenomenon, as the local parameters were not modified in these regions.

**Fig. 6.**
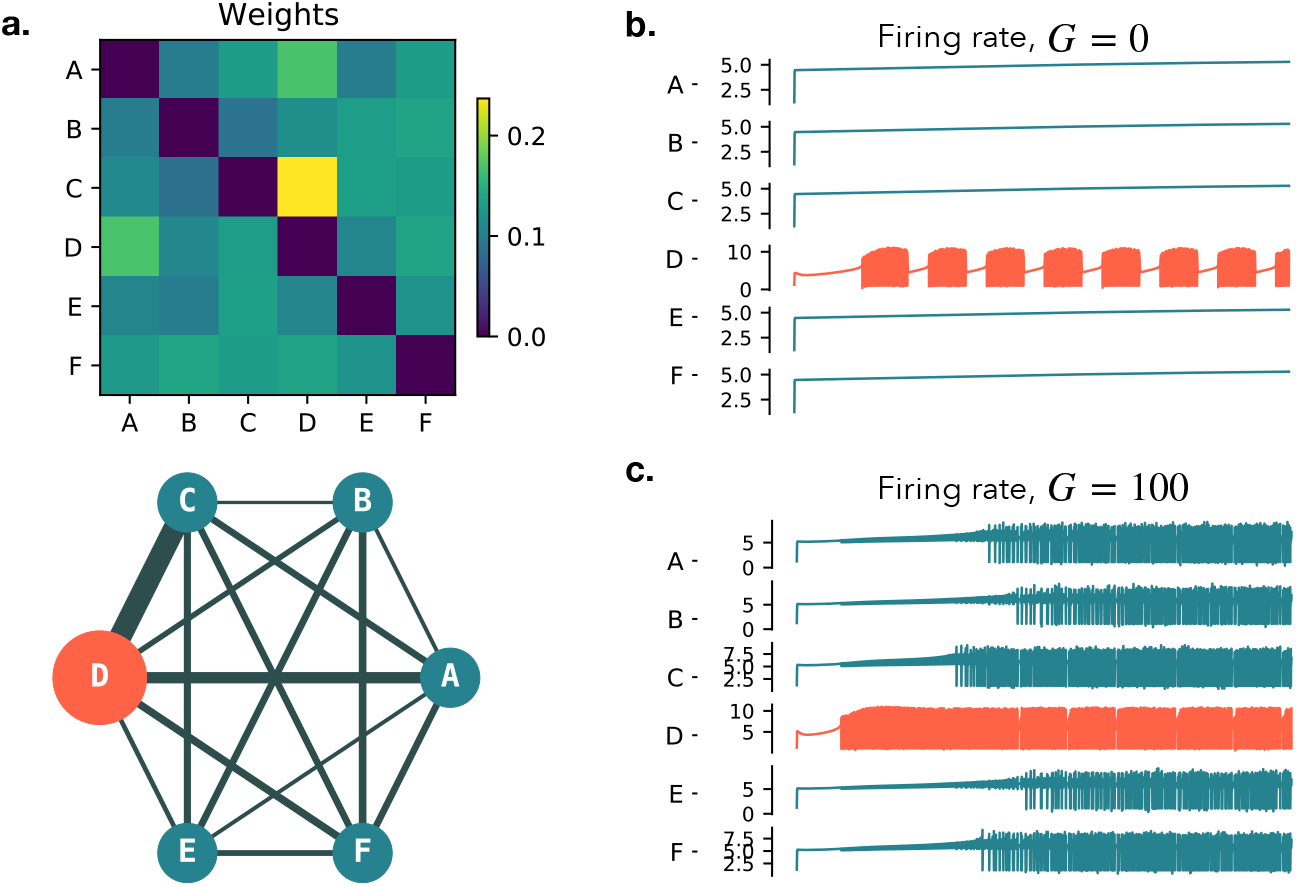
Network simulation of structurally connected neural mass models and propagation of pathological bursts: **(a)** Structural connectivity for six all-to-all connected nodes *A, B, C, D, E, F* with random weight allocation. Each node is described by a neural mass model derived as the mean-field approximation of a large population of HH type neurons. **(b)** When decoupled (Global Coupling *G* = 0), all the nodes operate in a ‘healthy’ regime with the potassium concentration in their bath set to low values [*K*^+^]_*bath*_ = 5.5, except for node *D* which is tuned into a pathological regime [*K*^+^]_*bath*_ = 15.5 characterized by the spontaneous presence of bursts. **(c)** When the global coupling is sufficiently increased, the pathological value of [*K*^+^]_*bath*_ in node *D* generates bursts that diffuse through the connectome mimicking the spreading of a seizure.

This simulation serves as a proof of concept to illustrate how local pathological activity can propagate through a network depending on the strength of coupling. We used a single representative realization of randomly weighted structural connectivity. While we did not perform a systematic exploration of different realizations or coupling strengths, we observed that the qualitative behavior — namely, the emergence of network-wide bursting beyond a critical coupling threshold — remains robust across similar setups. The model is compatible with empirical connectome data and can be readily extended to simulations using realistic brain network architectures.

### 1.6 Limitations of the model

The mean-field model derived in this work relies on approximations and heuristic arguments that, on the one hand, allow a closed-form derivation of the mean-field equations (38), and on the other hand restrict its validity to a limited regime of activity corresponding to quasi-synchronous neuronal populations. Therefore, rather than an exact mean-field representation, the model provides a qualitatively realistic description of a mesoscopic population of connected neurons driven by ion exchange dynamics. Moreover, the discrepancy between the two modalities would have likely been smaller if for the neural network we also adopted a gating variable that is mesoscopic and identical across the spiking neurons, as in similar works [49–51]. However, here we demonstrate the validity of the mean-field approximation even for the more natural, microscopic representation of the gating variable in the neural network.

The approximation of the membrane potential dynamics as a step-wise quadratic function and the assumption of Lorentzian distributed membrane potential in the population allowed us to apply the Ott-Antonsen Ansatz and to develop the mean-field approximation of a heterogeneous network of biophysical neurons driven by ion-exchange dynamics coupled all-to-all via conductance-based coupling. With these approximations, the mean-field equations do not show a one-to-one correspondence with the neural network simulations, except in regimes where the distribution of the membrane potentials of the neurons is well described by a Lorentzian (Fig. 3.a). In fact, it is not guaranteed that such a population of HH-type neurons allows for a closed formulation of the thermodynamic limit. For example, the distribution of the membrane potentials resembles a Lorentzian only in some cases (e.g., when all the neurons are unimodally distributed below or above the threshold; Fig. 4.b, bottom left), while it can deviate from it in other moments (e.g., when a subpopulation of neurons transition suprathreshold while another population remains subthreshold; Fig. 4.b, bottom right).

The mean-field model could be improved, for example by allowing for a bimodal distribution of the membrane potential (e.g., a double-Lorentzian approximation), although this condition is not guaranteed to allow a closed-form derivation of the mean-field equations. Furthermore, the parabola coefficients *c*_−_, *c*_+_, *R*_*−*_, *R*_+_ were fixed as constants, however, these coefficients could be made functions of the slow variables and the gating variable, which might unveil new dynamical regimes and extend the validity of the thermodynamic limit beyond the regimes described in this work. Also, in the case of constant values, an in-depth exploration of the parameter space is required to fully characterize the model and its bifurcation structure. Other limiting assumptions are the moment closure condition (19) and the assumptions that the functions (3) averaged across the neuronal population can be expressed as functions of the average membrane potential *V* and gating variable *n* (which is only true in the cases where the functions (3) can be reasonably approximated as linear functions in a range of *V* and *n*).

The definition of the firing rate in the thermodynamic limit was approximated assuming a firing threshold for membrane potential going to infinity as in (25), however, more refined definitions (e.g., accounting for a heterogeneous firing threshold) can be considered [82]. In this work, we showed that the model can retrieve emergent network behaviors similar to those observed in neuronal network simulations and in vitro. This correspondence is not quantitative because the network size is finite, and, in experimental observations (in vivo/in vitro networks), the connectivity profile is not all-to-all connected. Despite its simplicity, the model displays desired properties such as bistability in healthy and pathological regimes (Fig. 5) and sensitivity to external coupling (Fig. 6). This approach, taking into account key biophysical details, offers a first step in considering the role of the glia in neural tissue excitability. Following this direction, other ions, such as calcium should be taken into consideration, as well as other effects such as plastic synapses, random connectivity, noise, adaptation, spike-timing-dependent plasticity, as already discussed in the Introduction.

## Discussion

A quest of modern neuroscience is understanding and explaining how the brain operates for healthy individuals, and how it deviates from its healthy state in case of different diseases and disorders. Most brain imaging techniques register the collective electrical and metabolic fluctuations of large neuronal populations. Because these fluctuations emerge from the electrochemical interactions of many neural cells, they cannot be fully understood at the microscopic level (e.g., as the mechanism of a single neuron). Instead, they must be studied at a mesoscopic level, where the emergent behavior of the entire population can be observed. Also, besides the recent progress [83], it is still not possible to compute the behavior of a few billion such neurons (which comprises a mammalian brain), and even if it was, it remains debatable what knowledge we would gain from such a complex system [84]. Hence, an appropriate way to model recorded brain activities is to write mathematical descriptions for large neuronal populations [7], using formalism from statistical physics and mathematics. In this case, one can measure population properties such as the mean membrane potential, rather than the activity of an individual neuron. Some of these observables acquire meaning only in the context of mean-field analyses, for example, temperature or pressure in the case of molecular ensembles, order parameter [85] and mean ensemble frequency [86] for coupled oscillators, or more specifically the firing rate in the case of neuronal population [87]. However, to the best of our knowledge, no theory to date can explain the behavior of a large population of neurons (brain areas) from the perspective of the driving mechanism of the neuronal activities, that is the ion exchange and transportation dynamics at the cellular level. Different pathological trials and experiments have already revealed that changes in ion concentration in brain regions could result in different brain dysfunctions but the pathway of these phenomena is still unclear.

In this study, we developed a biophysically inspired mean-field model for a network of all-to-all connected, heterogeneous Hodgkin–Huxley type neurons, which are characterized by ion-exchange mechanism across the cellular space. Unlike phenomenological or reduced models, the Hodgkin–Huxley framework allows us to retain explicit ion exchange dynamics, which are essential for linking membrane behavior to extracellular potassium fluctuations. This level of biophysical detail is crucial for modeling pathological regimes such as seizure onset and propagation.

The intermediate approximation into the step-wise quadratic function allowed us to apply the analytical formalism to the complex Hodgkin–Huxley type equations. The further assumption that the membrane potentials of neurons in a large population are distributed according to a Lorentzian (Lorentzian ansatz) makes the mean-field approximation analytically tractable. From the simulation of the network behavior of such neurons, it is evident that the mean-field model captures the dynamic behavior of the network when the population is highly synchronized. This assumption might find applications in the study of seizure dynamics. Future work is needed to explore the non-synchronous cases.

The bifurcation analysis reveals that the mean-field model is capable of qualitatively capturing emergent neuronal network states observed in numerical simulations, as well as in vitro (e.g., Fig. 4). The significance of such qualitative analysis lies in its relevance to a recently proposed classification of seizure dynamotypes, which categorizes seizures based on their observable characteristics and dynamic composition [81]. This classification was supported by a model-based bifurcation analysis using the “Epileptor” model as a neural mass model [31]. Our model complements the Epileptor by allowing the interpretation of parameters in terms of measurable microscopic quantities, specifically ion concentrations, rather than phenomenological parameters. Therefore, our model allows us to translate the classification of seizure dynamics from electrophysiology recording with mesoscopic scale resolution (S/M/EEG) to experiments with microscopic details, as we have shown invitro. While our study contributes to both the qualitative agreement between our model and in-vitro seizure patterns and the improved interpretability of phenomenological models, a full bifurcation analysis, beyond the one presented in this work is required to align our results with the dynamical patterns of the Epileptor model.

The derived mean-field model relates the slow-timescale biophysical mechanism of ion exchange and transportation in the brain to the fast-timescale electrical activities of large neuronal ensembles. In fact, as demonstrated via brain imaging of different modalities [88], ionic regulation acts on a relatively slow time scale on the fast electrical activity of neurons.

Analyzing the model, we found the potassium concentration in the external bath (parameter [*K*^+^]_*bath*_) to be a significant determinant of the dynamics, monitoring the biophysical state of the neuronal population (i.e., a brain region). For low ion concentrations in the external bath, the mean-field model demonstrates the existence of healthy brain dynamics. Increasing the excitability in terms of the ion concentration leads to the appearance of spike trains, tonic spiking, bursting, seizure-like events, status epilepticus-like events, and depolarization block, similar to the Epileptor model [31, 89, 90].

For several values of the potassium concentration in the external bath, or given an external input current, we showed that our model can be tuned into a bistable regime characterized by the coexistence of high and low firing rate states, which is the hallmark of many mean-field representations of linear [91, 92] or quadratic integrate and fire neurons [25, 93], and rate models [22]. Bistability is a desired feature for mean-field models. For example, bistability might provide a mechanism for the dynamic occurrence of so-called up (high firing) and down (low firing) states, as observed both in vitro [94, 95] and in vivo under several conditions such as quiet waking, anesthesia, slow-wave sleep and during perceptual task across several species [96–100]. In fact, when bistable neural masses are coupled through a connectome and driven by a fine-tuned stochastic input noise, the simulated activities in brain regions can spontaneously jump between high and low firing rate states, which provides a mechanism for dynamic Functional Connectivity [10], as observed in large-scale brain recordings [101].

Thus, the derived mean-field model links the presence of high and low firing rate states as well as the spiking and bursting neuroelectric behaviors with the biophysical state of the neuronal ensemble (brain regions) in terms of the ion concentrations across the cellular space. The effect of constant stimulus current is also analyzed and it is observed that even within the healthy regime, several stimulations, which could either correspond to external stimuli or inputs from some other brain regions, could generate transient spiking and bursting activity (Fig. 5.d). This result is particularly interesting in the case of brain stimulation. For example, several kinds of brain stimulation protocols in epileptic patients have already been found to generate epileptic seizures in pathological practice [102, 103]. Our results demonstrate that these phenomena could be reproduced and tracked in a biophysical-inspired modeling approach. Therefore, we assume that the derived model could potentially be applied to improve predictive capacities in several types of brain disorders, particularly epilepsy.

Using conductance-based coupling between six neuronal masses we also demonstrated that a hyper-excited population of neurons can propagate bursting and spiking behavior to otherwise healthy populations. This could lead the path to understanding how brain signals propagate as a coordinated phenomenon depending on the distribution of biophysical quantities and structural as well as architectural heterogeneity on the complex network structure of the connectome. Until now, mean-field models used in large-scale network simulations, like the Virtual Brain [7], did not take into consideration the extracellular space. Our mean-field model is a first step towards the integration of biophysical processes that may play a key role in controlling network behavior, as shown at the spiking network level [104].

In this work, we focused on [*K*^+^], given its known role in neuronal activity. Changes in *K*^+^ occur when the brain alternates between arousal and sleep [61]. Although a causal relationship is not clearly established, seizures and spreading depression, which is assumed to underlie certain forms of migraine [105, 106], are associated with large (*>*6 mM) and very large (*>*12 mM) concentrations of *K*^+^ [105, 107]. The invariant increase of *K*^+^ concentration during seizure [108], provokes a saturation of potassium buffering by the astrocytes. In fact, the extracellular concentration of *K*^+^ is tightly controlled by astrocytes [73, 74, 76]. Most large-scale simulations do not integrate glial cells, which make up half of the brain cells [73, 76]. Their functions are altered in most, if not all, brain disorders, in particular, epilepsy [75, 77]. Our approach indirectly accounts for the astrocytic control of extracellular *K*^+^. Other ion species also vary during arousal/sleep and seizures, in particular, *Ca*^2+^ [61, 109–111]. A decrease in *Ca*^2+^ will decrease neurotransmission and thus change cell-to-cell communication [109–111]. Future studies are needed to integrate *Ca*^2+^ in neural mass models. This paper serves as an introductory exploration of our novel approach, aiming to establish its feasibility and potential applications for modeling large ensembles of biophysically realistic neurons. A comprehensive and systematic investigation to address the limitations discussed in Section ‘Limitations of the model’ and FigS1 will be undertaken in future studies.

To date, the Epileptor model [31] is a gold standard for inferring the location of the epileptogenic zone, with direct applications in clinics [112, 113]. Nonetheless, the Epileptor model remains phenomenological, and parameters (e.g., epileptogenicity) are non-interpretable in terms of measurable quantities. By addressing the biophysical information at the neuronal level, our mean-field formalism allows keeping biophysical interpretability while bridging between micro-to macro-scale mechanisms. The mean-field model derived in this work aggregates a large class of brain activities and repertoire of patterns into a single neural mass model, with direct correspondence to biophysically relevant parameters. This approach could serve as a computational baseline to address core questions in epilepsy research. In particular, how to identify the multiscale mechanisms implicated in epileptogenicity and propagation of seizures. This would eventually lead to establishing pathologically measurable bio-markers for large-scale brain activities and consequently offering therapeutic targets for different brain dysfunctions.

## 2 Materials and Methods

### 2.1 Immature hippocampus in vitro preparation

All protocols were designed and approved according to INSERM and international guidelines for experimental animal care and use. Experiments were performed on intact hippocampi taken from FVB NG mice between postnatal (P) days 5 and 7 (P0 was the day of birth). The immature mice were decapitated rapidly, and the brain was extracted from the skull and transferred to oxygenated (95% O_2_ / 5% CO_2_) ice cold (4 °C) artificial cerebrospinal fluid (aCSF) containing (in mM): NaCl 126; KCl 3.5; CaCl_2_ 2.0; MgCl_2_ 1.3; NaHCO_3_ 25; NaHPO_4_ 1.2; glucose 10 (pH = 7.3). After at least 2 hours of incubation at room temperature in aCSF, hippocampi were transferred to the recording chamber. To ensure a high perfusion rate (15 ml/min) we used a closed-circuit perfusion system with recycling driven by a peristaltic pump: 250 ml of solution was used per hippocampus and per condition. The pH (7.3) and temperature (33 °C) were controlled during all experiments. After 30 min baseline recording in Mg^2+^ containing aCSF solution, the media was switched to one without added Mg^2+^ ion. In this condition, the extracellular concentration of Mg^2+^ may be influenced by other constituents of the aCSF, possibly near 0.08 mM [114]. Therefore, we use the term low-Mg^2+^ aCSF rather than zero-Mg^2+^ aCSF.

### 2.2 Magnesium removal as a mean to influence the external potassium dynamics

The model of epileptic discharges presented in our study was first introduced over 20 years ago [115] and has since become a well-established paradigm for screening potential antiepileptic drugs and research on the mechanism of epileptic seizure [116].

The membrane of hippocampal neurons is equipped with N-methyl-D-aspartate type glutamate receptors (NMDARs). These receptors have a very high affinity for glutamate and can, in principle, be activated by ambient glutamate present at low concentrations in the brain extracellular fluid (ECF). Under normal physiological conditions, this activation does not occur because extracellular magnesium ions (*Mg*^2+^) block the NMDAR channel at membrane potentials more negative than about –50 mV; this voltage-dependent block prevents receptor activation at rest. When extracellular magnesium is removed, the block is relieved, allowing NMDARs to be activated, leading to neuronal depolarization toward the action potential threshold [117]. In addition, as a divalent cation, *Mg*^2+^ interacts with the negatively charged neuronal membrane, contributing to the stabilization of the resting membrane potential. Lowering extracellular magnesium concentration disrupts this effect, resulting in membrane depolarization [118].

Consequently, magnesium removal not only facilitates NMDAR-dependent depolarization, but also directly depolarizes neurons. This depolarization increases the driving force for outward potassium currents through *K*^+^ channels, meaning that variations in *Mg*^2+^ can indirectly influence external potassium dynamics during neuronal activity.

### 2.3 Electrical activity monitoring

Extracellular glass electrode filled with low Mg^2+^ aCSF was placed into the mid CA1 region of the ventral hippocampus. Local field potentials (LFPs) were amplified with a MultiClamp700B (Molecular Devices, San Jose, CA) amplifier for DC-coupled recording, then digitized with a Digidata 1440 (Molecular Devices, San Jose, CA), stored on the hard drive of the personal computer and displayed using PClamp 9 software (Molecular Devices, San Jose, CA).

### 2.4 External Potassium concentration Measurements

Potassium-selective microelectrodes were prepared using the method described by Heinemann and Arens [119]. In brief, electrodes were pulled from double-barrel theta glass (TG150-4, Warner Instruments, Hamden, CT, USA). The reference barrel was filled with 154 mM NaCl solution. The silanized ion-sensitive barrel tip (5% trimethyl-1-chlorosilane in dichloromethane)was filled with a potassium ionophore I cocktail A (60031 Fluka distributed by Sigma-Aldrich, Lyon, France) and backfilled with 100 mM KCl. Measurements of [K^+^]_ext_-dependent potentials were performed using a high-impedance differential DC amplifier (kindly provided by Dr. U Heinemann [119]) equipped with negative capacitance feedback control, which permitted recordings of relatively rapid changes in [K^+^]_ext_ (time constants 50–200 ms). The electrodes were calibrated before each experiment and had a sensitivity of 42-63 mV/mM. The tip of the electrode was placed into the CA1 region of the ventral hippocampus at 200-300µm distance from the LFP recording electrode at 150-200 µm depth that, at P5-P6 age, corresponds to the Stratum Radiatum.

### 2.5 Biophysical neuron model

The membrane potential of a single neuron in the brain is related to an ion-exchange mechanism in intracellular and extracellular space. The concentrations of potassium, sodium, and chloride in the intracellular and extracellular space along with the active transport pump (*Na*^+^/*K*^+^ pump) in the cell membrane of neurons generate input currents that drive the electrical behavior in terms of the evolution of its membrane potential. The ion-exchange mechanisms in the cellular microenvironment, including local diffusion, glial buffering, ion pumps, and ion channels, have been mathematically modeled based on conductance-based ion dynamics to reflect the ‘healthy’ and seizure behaviors in single neurons [68, 78, 79, 120]. Moreover, conductance-based couplings between the spiking neurons have been already implemented in neural mass models [58, 59, 91, 93, 121], but without an extracellular exchange mechanism. The mechanisms of ion exchange in the intracellular and extracellular space of the neuronal membrane are represented schematically in Fig. 1.

This biophysical interaction and ion-exchange mechanism across the membrane of a neuronal cell can be described as a Hodgkin–Huxley type dynamical process, represented by the following dynamical system as described in [78].

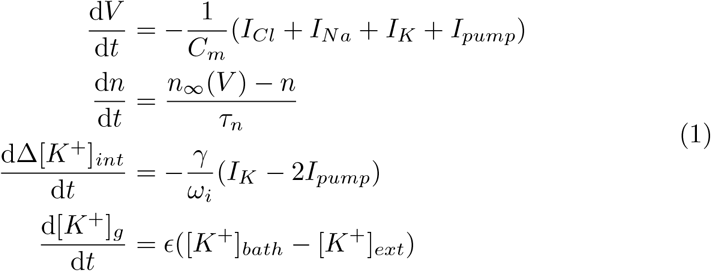

This model represents the ion-exchange mechanism of a single conductance-based neuron in terms of membrane potential *V*, the potassium conductance gating variable *n*, intracellular potassium concentration variation Δ[*K*^+^]_*int*_, and extracellular potassium buffering by the external bath [*K*^+^]_*g*_. This mechanism considers ion exchange through the chloride, sodium, potassium, voltage-gated channels, intracellular sodium, and extracellular potassium concentration gradients and leak currents. The intrinsic ionic currents, the sodium-potassium pump current, and potassium diffusion regulate the different ion concentrations. The Nernst equation was used to couple the neuron’s membrane potential with the concentrations of the ionic currents. This mechanism gives rise to a slow-fast dynamical system in which the membrane potential *V* and potassium conductance gating variable *n* constitute the fast subsystem and the slow subsystem is represented in terms of the variation in the intracellular potassium concentration Δ[*K*^+^]_*int*_ and extracellular potassium buffering by the external bath [*K*^+^]_*g*_ (Fig. 2.b). In Eq. (1) the input currents due to different ionic substances and pump are represented as follows [68, 78]:

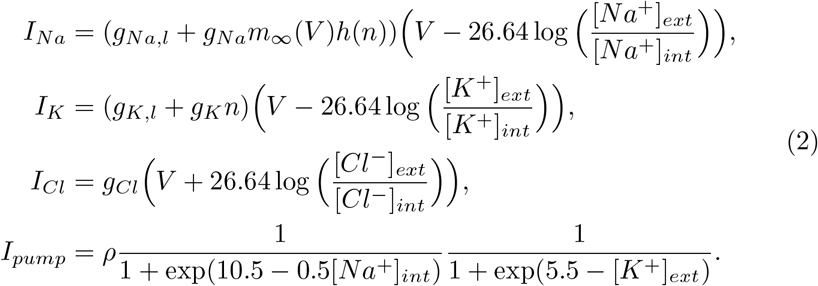

The gating functions for the conductance are modeled as

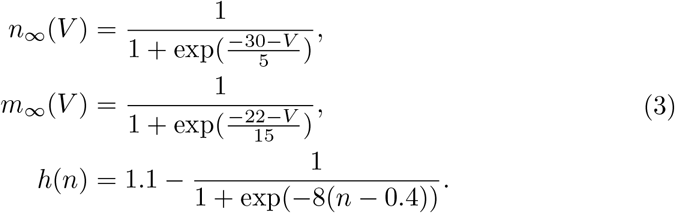

Notice that, compared to [78], these equations were reparametrized to account for changes in timescales (see also 1). In this model the concentration of chloride ions inside and outside of the membrane is invariant. i.e., [*Cl*^−^]_*ext*_ = [*Cl*^−^]_0,*ext*_ and [*Cl*^−^]_*int*_ = [*Cl*^−^]_0,*int*_. The extracellular and intracellular concentrations of sodium and potassium ions are represented in terms of the state variables Δ[*K*^+^]_*int*_, [*K*^+^]_*g*_ as follows [78]:

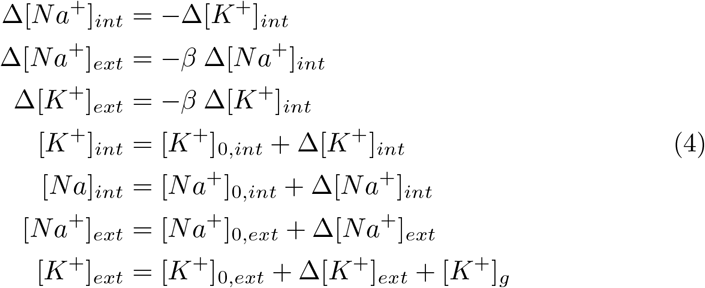

The biophysically relevant values of the parameters could be obtained from several previous studies, both from in vivo and in vitro experiments [68]. The parameters used for the single-neuron simulations are shown in Table-1, and allow the reproduction of several bursting and oscillatory regimes by varying the parameter [*K*^+^]_*bath*_ (Fig. 2.a).

### 2.6 Network of coupled biophysical neurons

We aim to develop a mean-field model for a heterogeneous population of the all-to-all coupled biophysical neurons described by Eq. (1). Our strategy consists of using a series of approximations and heuristic arguments to obtain a putative functional form for the mean-field model, which is capable of simulating the mean activity of a large population of Hodgkin–Huxley (HH) type neurons operating near synchrony. The basic mechanism of synaptic transmission can be described by the arrival of a spike at the presynaptic terminal, resulting in an influx of calcium which depolarizes the presynaptic terminals and thus stimulates the release of neurotransmitters into the synaptic cleft. The neurotransmitters play a crucial role in opening the postsynaptic channels, which causes the flow of ionic currents across the membrane. This flow creates an (inhibitory or excitatory) post-synaptic potential in the efferent neuron. We take this into account by adding a synaptic input current in the membrane potential equation at neuron *i* using a conductance-based model

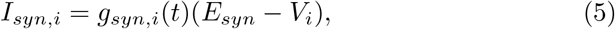

where *g*_*syn,i*_ is the synaptic conductance, *V*_*i*_ is the postsynaptic potential and *E*_*syn*_ is the synaptic reversal potential at which the direction of the net current flow reverses. In the present work, we fixed *E*_*syn*_ = 0mV, corresponding to excitatory neurons. Lower reversal potential values (e.g., *E*_*syn*_ = −80mV) model inhibitory neurons. Typically, at the arrival time 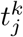 of the *k*-th spike from neuron *j*, the synaptic conductance quickly rises and consequently undergoes an exponential decay with some rate constant. This mechanism is captured by defining the conductance dynamics *g*_*syn,i*_ ∝ *Js*_*i*_(*t*) where *J* is a constant conductance value and the adimensional dynamical variable *s*_*i*_(*t*) is the solution of the following equation (see for example [28]):

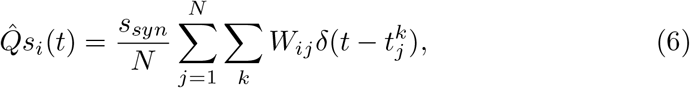

where *s*_*syn*_ = 1 is the synaptic strength, *W*_*ij*_ is the synaptic weight which here we assume to be equal to 1 for all neurons *i* and *j*, and the operator 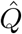 can be defined with various levels of accuracy as

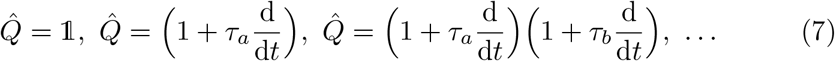

Here, for simplicity, we select the first model, which leads to

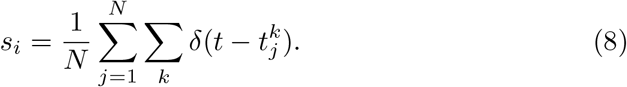

Under the assumption of instantaneous synaptic transmission and homogeneous all-to-all coupling, the synaptic activation variable *s*_*i*_ is the same for all neurons and corresponds to the population firing rate, which we denote by *r*.

In the coupled system, the membrane potential equation for neuron *i* is defined by:

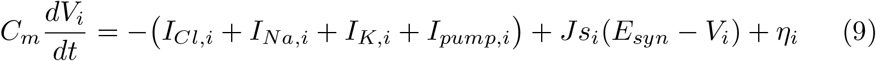

where we fix the capacitance to *C*_*m*_ = 1*n*F, and the term *η*_*i*_ represents a heterogeneous noise current distributed according to a Lorentzian distribution with half-width Δ and location of the center at 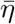

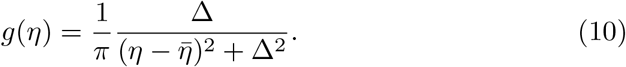

### 2.7 Continuity equation

In the continuous formulation, in the limit of large population *N* → ∞, the density of neurons in a phase space point (*V, n*, Δ [*K*^+^]_int_, [*K*^+^]_*g*_) at time *t* and excitability *η* is described by the population density function *ρ*(*t, V, n*, Δ [*K*^+^]_int_, [*K*^+^]_*g*_, *η, t*), and the continuity equation holds

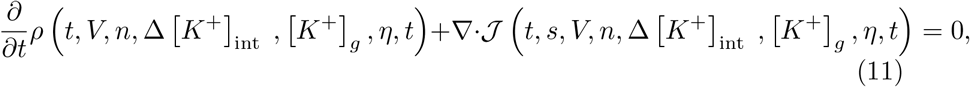

where 𝒥 is the flux along the *V, n*, Δ [*K*^+^]_int_, and [*K*^+^]_*g*_ directions. Since our system displays a fast and a slow subsystem (e.g., Fig. 2.b), we treat the variables Δ [*K*^+^]_int_, and [*K*^+^]_*g*_ as slowly varying parameters. In other words, we assume they are nearly constant over the timescale of the fast variables *V* and *n*. These slow variables are in addition considered to be mesoscopic, meaning they are identical for every neuron in the population. In this setup, the slow variables Δ [*K*^+^]_int_, and [*K*^+^]_*g*_ determine the long-term behavior of the system, while the equations for *V*, and *n* describe the rapid responses around this slow evolution. Furthermore, we consider that the potassium concentrations are homogeneous across the neuronal population, and we assume this holds also for the other ion concentrations.

Therefore, we write the population density function for the fast subsystem as

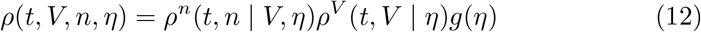

which was here expressed in the conditional form independent of the potassium variables. The continuity equation reads

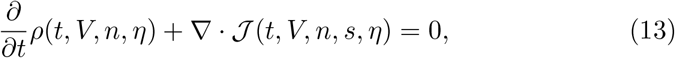

where the flux in the *V* and *n* directions is

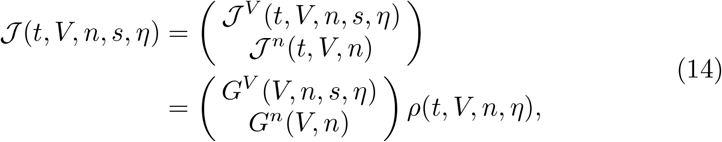

With

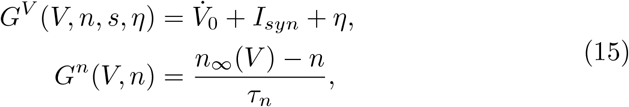

And

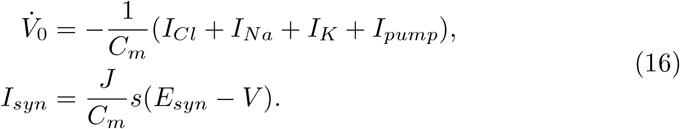

We note that, under the assumption of globally shared gating and ion concentration variables across the neuronal population, the resulting mean-field equations can also be derived using simpler methods as proposed by Guerriero et al. [58]. In this work, we follow the more general formalism of Chen and Campbell [59], which makes the role of key approximations (e.g., moment closure, vanishing flux at boundaries) explicit. This also facilitates potential generalizations to settings with partial heterogeneity or dynamic gating distributions.

Following [59], we obtain a modified version of the continuity equation by integrating the continuity equation 13 with respect to *n*. Using the normalization condition on the marginal density of *n*, the first term gives

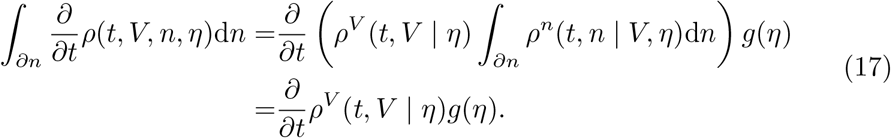

Similarly, integrating the second term in the continuity equation 13 with respect to *n* we obtain

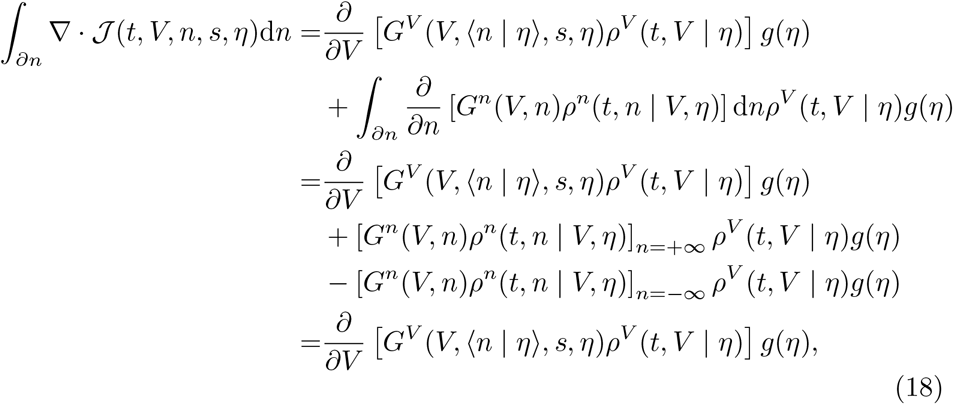

where we assumed that the flux along *n* vanishes on the boundary *∂n*. Next we assume a first-order moment closure condition for the variable *n* [59], justified by the numerical simulations of the full network (see Fig. S2) which show that for most of the neurons (close to 99 % for the value of Δ same as in the other simulations) the mean of the population is well capturing the behavior of the single neurons [122]. Finally, putting together these factors and assuming that *n* can be treated as a collective variable for each neuron (see *Limitations of the model* section) we arrive to

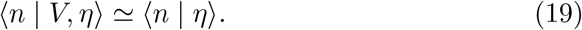

From here we obtain a modified version of the continuity equation for the fast subsystem

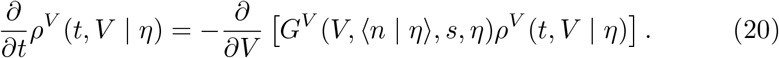

The validity of the first moment closure, Eqs. (19), as in [59], is supported by the numerical simulations, which show that, both, during the silent regime and when seizure-like events occur, *n*_*i*_ for most neurons track the network averaged ⟨*n* |*V, η*⟩ . In particular, it is less than 2 % of the neurons that fire while the mean is low, and vice-versa, Fig. S2. In less synchronized scenarios (larger Δ or smaller *J*), however, this value would increase, but the mean would always capture the qualitative behaviour of the population.

### 2.8 Stepwise quadratic approximation

For a decoupled system described by Eqs. (1), considering the potassium variables as slowly varying parameters, and for any value of the gating variable *n*, the equation for the membrane potential 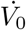 (16) has the profile of a cubic-like function (Fig. 2.c). Here, we approximate this function as:

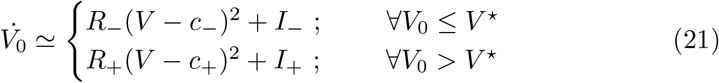

corresponding to two parabolas with opposite curvature (*R*_−_ > 0, *R*_+_ < 0), centered at *c*_−_ and *c*_+_ and shifted by *I*_+_ and *I*_−_, respectively (Fig. 2.d). The parameter *V* ^⋆^ defines the intersection point of the two parabolas. In general, the coefficients of the parabolas (*c*_±_, *I*_±_, *R*_±_) would be functions of *n*(but also of Δ[*K*^+^]_*int*_ and [*K*^+^]_*g*_). In the expression (20), this dependence is reduced to *c*_±_ = *c*_±_(⟨*n* |*η* ⟩), *I*_±_ = *I*_±_(⟨*n*| *η*⟩), and *R*_±_ = *R*_±_(⟨*n* |*η* ⟩). Here, to further simplify our problem, we consider these as free parameters, that will appear in the mean-field model, and that can be tuned to fit the average activity of a large neuronal network (see *Limitations of the model* section).

### 2.9 Steady-state solution and Lorentzian Ansatz

The steady-state solution for the (20) corresponds to

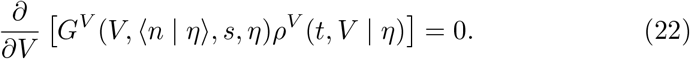

In standard mean-field reductions (e.g., [25, 59]), the function *G*^*V*^ is quadratic in *V*, with two fixed points, one of which is stable, and the steady-state solution for *ρ*^*V*^ (*t, V*| *η*) is the inverse of a quadratic i.e., a Lorentzian distribution. The Lorentzian Ansatz assumes that, at all times, such a distribution is preserved during the system’s dynamics. These conditions allow the derivation of closed solutions for a system of coupled spiking neurons. In our case, where *G*^*V*^ has either one or three zeros, the steady state will correspond to delta functions at the fixed points, which are either one (stable) or three (of which two are stable). Therefore, a double-Lorentzian (or a piece-wise Lorenzian) could be a suitable form for *ρ*^*V*^ (*t, V* |*η*). However, it is not clear under which conditions such an assumption would allow a solution to the continuity equation (20).

Nonetheless, in special cases, such as when the neuron population is acting close to synchrony, the neurons’ membrane potential distribution is faithfully described by a single Lorentzian peaked below or above the threshold (e.g., Fig.3.a; notice that the orange distribution is bimodal, as a few neurons are idling in the subthreshold state). To proceed with the model derivation, we assume that the distribution of membrane potentials for neurons with excitability *η* is described by a single Lorentzian distribution (Fig. 2.e) on either side of the threshold *V* ^⋆^

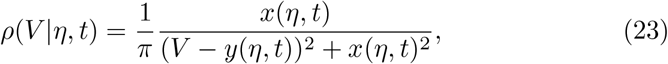

where *x*(*η, t*) and *y*(*η, t*) represent the width and the mean of the distribution, respectively. Thus, the validity of this approximation is limited to the cases when the neuronal population is distributed unimodally. Other cases, such as when a transition from sub-to suprathreshold activity of a subpopulation is described by the formation of two populations (e.g., Suppl. Fig.2.b), are not captured unless such transition is rapid enough. With these assumptions, we will derive the mean-field model equations in the next sections.

### 2.10 Firing rate definition

Previous mean-field derivations have adopted neuron models described by discontinuous quadratic equations, where the neuron is firing at a threshold *V*_*th*_, after which the voltage is reset to *V*_*reset*_. In these models, the threshold and reset are taken in the limit *V*_*th*_ = ™*V*_*reset*_ → ∞, and the quadratic expression for the membrane potential dynamics guarantees that a neuron takes a finite time to reach the threshold [25]. Therefore, the firing rate can be defined as the flux at infinity i.e.,

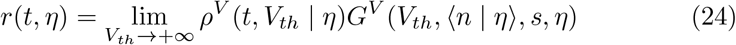

In our continuous neuron model, the spiking and resetting are built into the dynamic equation and can be driven by several mechanisms, including slow potassium dynamics. Let us consider the case of a steady input current affecting the membrane potential, such that the nullcline function (here described by the piece-wise quadratic function) will have either one zero (stable fixed point) or three zeros (two stable and one unstable). In this case, we can describe the firing at the pitchfork bifurcation, where the system transitions from three fixed points to one. At the bifurcation point, a neuron with membrane potential values at −∞ will be first driven by the positive quadratic trend, until it crosses the *V* ^⋆^ value, after which the negative parabolic trend will bring the neuron onto the fixed point. To have an operational definition of the firing rate and simplify our problem, we consider that once the *V* ^⋆^ value is crossed the neuron membrane potential will reach the peak value at +∞, driven by the positive quadratic trend. Of course, in our model, this value is never reached in practice, because–for increasing values of the membrane potential–the negative parabola brings the membrane potential back with increasing speed. In this approximation, the firing rate definition reduces to (see *Limitations of the model* section)

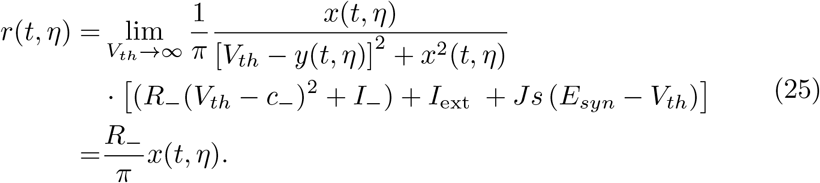

### 2.11 Mean-field variables

We describe the mean field variables as

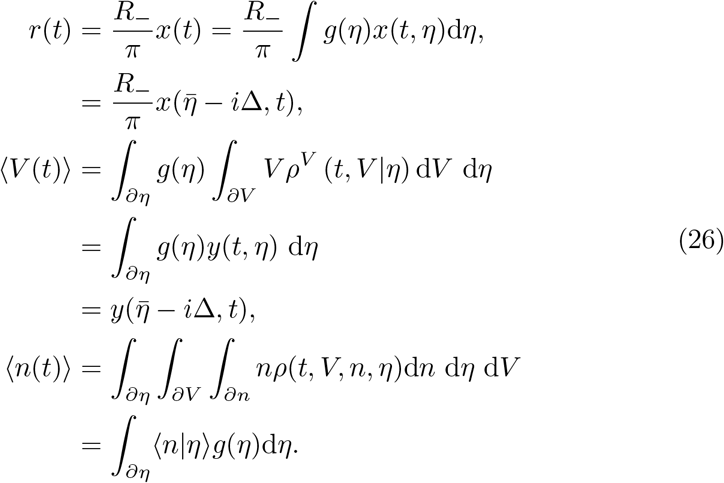

where we used the residue theorem to integrate out the parameter *η* (considering that there is only one pole of *g*(*η*) in the complex *η*-plane, we obtain that integrating out *η* corresponds to substituting 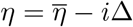).

Unlike the mean membrane potential *V* and the firing rate *r*, which can be explicitly derived from the continuity equation under the Lorentzian assumption, the expression for ⟨*n*(*t*) ⟩ in Eq. (26) is formal. In our mean-field model, the gating variable *n* is treated as a global population variable, evolving deterministically as a function of the average membrane potential. Therefore, ⟨*n*(*t*)⟩ corresponds to the collective gating variable assumed to be shared by all neurons, and is not computed by averaging distinct microscopic *n*_*i*_ values.

### 2.12 Mean-field dynamics for the gating variable

Following [59] we approximate the time derivative

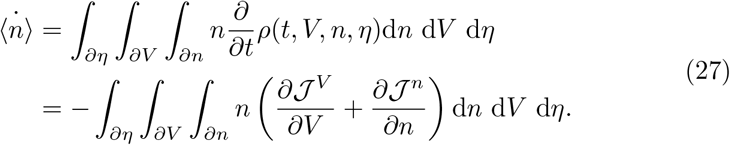

The first term is integrated by parts, yielding

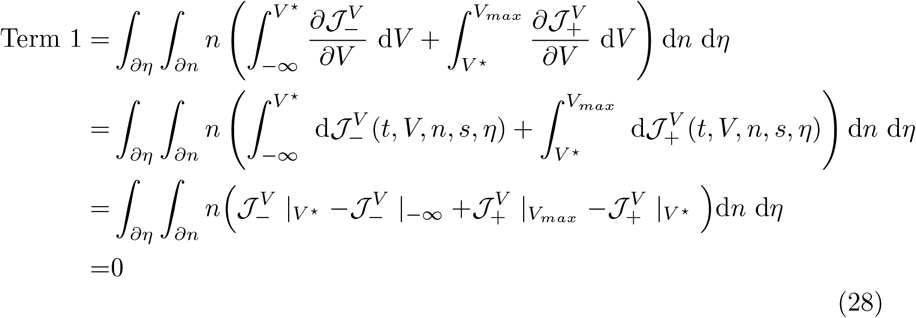

Where 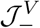 and 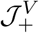 indicate the flux for *V* ≤ *V* ^⋆^ and *V* ≥ *V* ^⋆^ (preserved at *V*^⋆^, and governed by the positive and negative parabolas (21)), respectively, and where we assumed that the flux at *V*→ − ∞ and *V* →*V*_*max*_ is zero (a safe assumption, as a neuron is pushed back toward *V* ^⋆^ with (quadratic) infinite speed at these extreme values). The second term gives

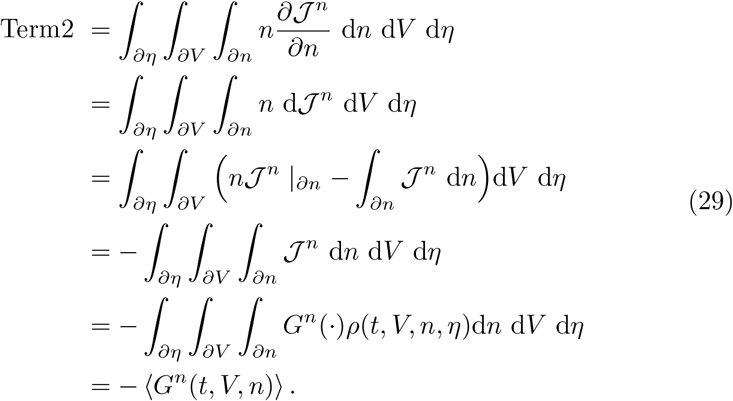

Imposing ⟨*n*_∞_(*V*) ⟩= *n*_∞_(⟨ *V* ⟩), we obtain the approximated result (see *Limitations of the model* section)

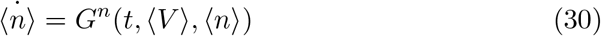

### 2.13 Derivation of mean-field equations

Starting from equations (21) and (5) the membrane potential dynamics for a coupled system of neurons is described on either side of *V* ^⋆^ by the following equation:

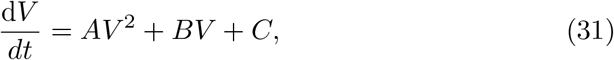

with

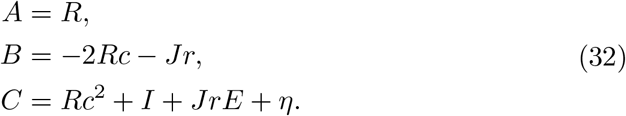

The parameters (*R, c, I*) were introduced in the previous sections as the curvature, center, and shift of the parabolas in the step-wise approximation. Here the indices (−, +) were dropped.

Substituting equations (31) and (23) in the continuity equation (20) we obtain a polynomial condition of the form

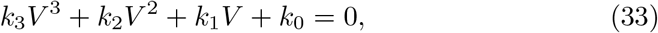

Since this condition must be satisfied for all values of *V*, the solution for *x*(*η, t*) and *y*(*η, t*) is obtained by imposing *k*_*I*_ ≡ 0, ∀*I* = 0, 1, 2, 3, leading to the mean-field equations

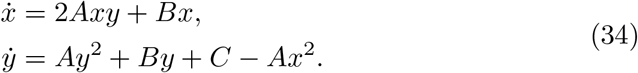

Defining *ω*(*η, t*) = *x*(*η, t*) + *iy*(*η, t*), we can recast the above equations in the complex form

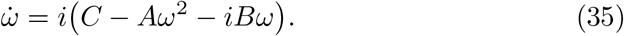

The mean-field equations are derived by integrating out *η* from the equation above. According to (26), we find 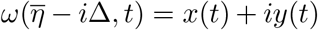, that substituted in (35) gives

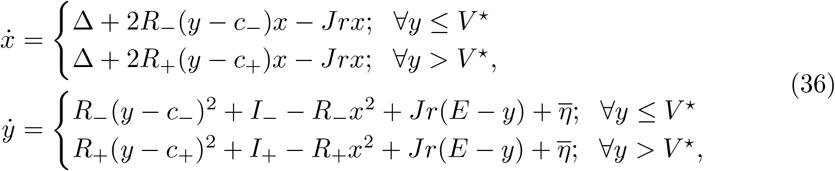

which is a reduction of a system of infinite all-to-all coupled neurons described by a step-wise quadratic equation.

### 2.14 Neural Mass Model equations

From equations (23) and (26), 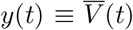 represents the average membrane potential of the population (from here on we will drop the ⟨·⟩ notation), while *x*(*t*) is an auxiliary variable, related to the firing rate as *r*(*t*) = *R*_−_*x*(*t*)/*π* (see previous section *Mean-field variables*). Finally, starting from equation (36), we reverse the stepwise quadratic approximation and reintroduce the original current-based formulation (Eq. (1)) into the mean-field model, thereby recovering the full dynamics of the slow variables Δ[*K*^+^]_*int*_, [*K*]_*g*_. The mean-field approximation for a population of HH-type neurons consists of a five-dimensional system:

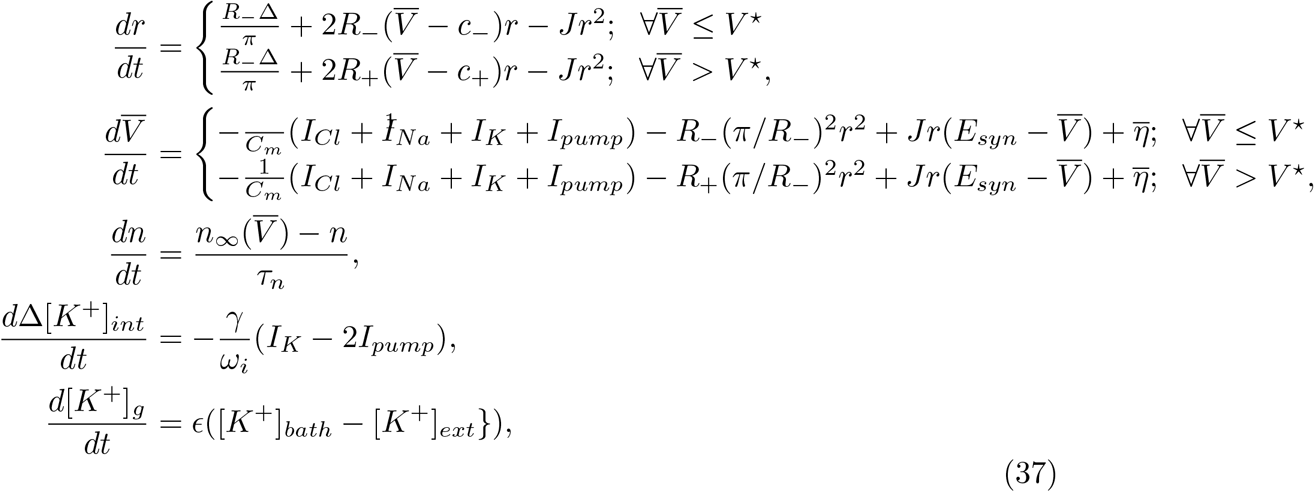

that we write in the compact form

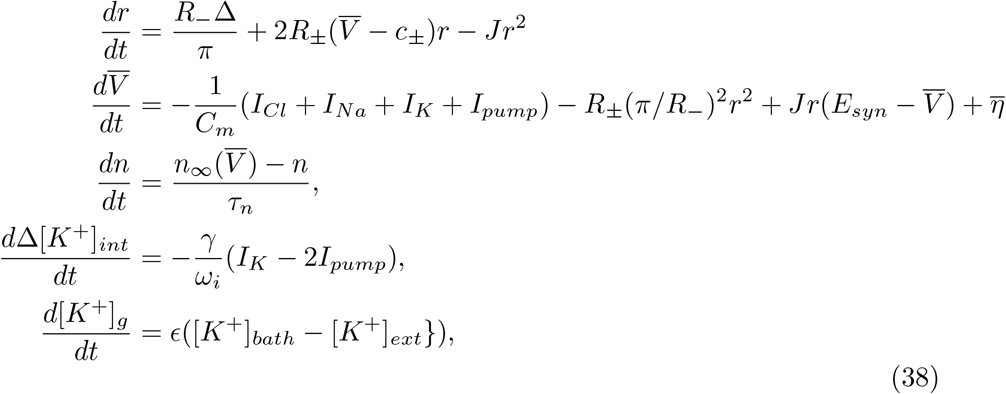

where the ± symbol refers to the cases 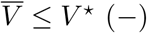and 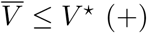.

### 2.15 Stimulation and Coupling of Neural Mass Models

In this work, the stimulation of the population is modeled by a common component *I*_*input*_(*t*) added to the membrane potential in Eq. 38 as:

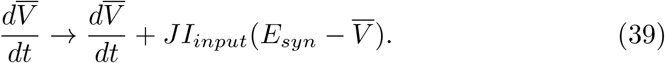

Accordingly, *N* populations described by Eq. 38 can be coupled together via their firing rate activity, so that the membrane potential equation at population *P* becomes:

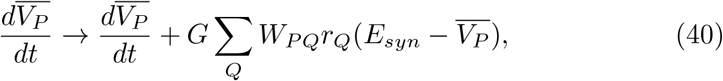

where *W*_*PQ*_ is the structural weight proportional to the white matter fiber tract density between populations *P* and *Q*, and *G* is the global coupling tuning the impact of the macroscopic network structure over the local mesoscopic dynamics.

The dynamics of a single neuron and single mean-field population were simulated in *Python*. Spiking neural network simulations were performed using *Brian2* package. Connectome-based simulations were run using The Virtual Brain platform [7]. In Table S2, we detail all the parameters and the values of the initial conditions used in the reported results.

## 3 Acknowledgments

## Funding

This research was supported by the European Union’s Horizon 2020 research and innovation program under Specific Grant Agreement 945539 (SGA3) Human Brain Project, No. 826421 Virtual Brain Cloud and No. 101147319 (EBRAINS 2.0 Project), and government grant managed by the Agence Nationale de la Recherche reference ANR-22-PESN-0012 (France 2030 program). CB received support from the Agence Nationale de la recherche projects ANR-17-CE37-0001-01 and ANR-20-NEUC-0005-02. GR is supported by the Marie Sklódowska–Curie Postdoctoral Fellowship (Project CAERUS) under the European Union’s Horizon Europe research and innovation programme, grant agreement No. 101199894. MD is supported by the Basque Government through the BERC 2022-2025 program and by the Ministry of Science and Innovation: BCAM Severo Ochoa accreditation CEX2021-001142-S / MICIU / AEI / 10.13039/501100011033.

## Author contributions

A.B, G.R., S.P., and V.K.J. designed research; G.R., A.B., C.C., K.G., D.D., L.MT, ML.S., M.D., S.P., and V.K.J. performed research; G.R., A.B., C.C., K.G., D.D., L.MT, M.D. simulated the models and analyzed the results; G.R., L.MT designed the figures. A.I. performed the experiments. G.R., A.B., D.D., ML. S., A.I., C.B., ML.L., S.P., and V.K.J. wrote the paper. *G.R. and *A.B. contributed equally to this work as first authors. +S.P. and +V.K.J. contributed equally to this work as the last authors. The authors declare no competing financial interests. Correspondence should be addressed to Giovanni Rabuffo at or Spase Petkoski at or Viktor K. Jirsa at.

## Competing interests

Authors declare that they have no competing interests.

## Data and materials availability

All data, code, and materials used in the analyses will be made available at a Github repository depending on the outcome of the editorial process.

## Supplementary information for

**Fig. S1.**
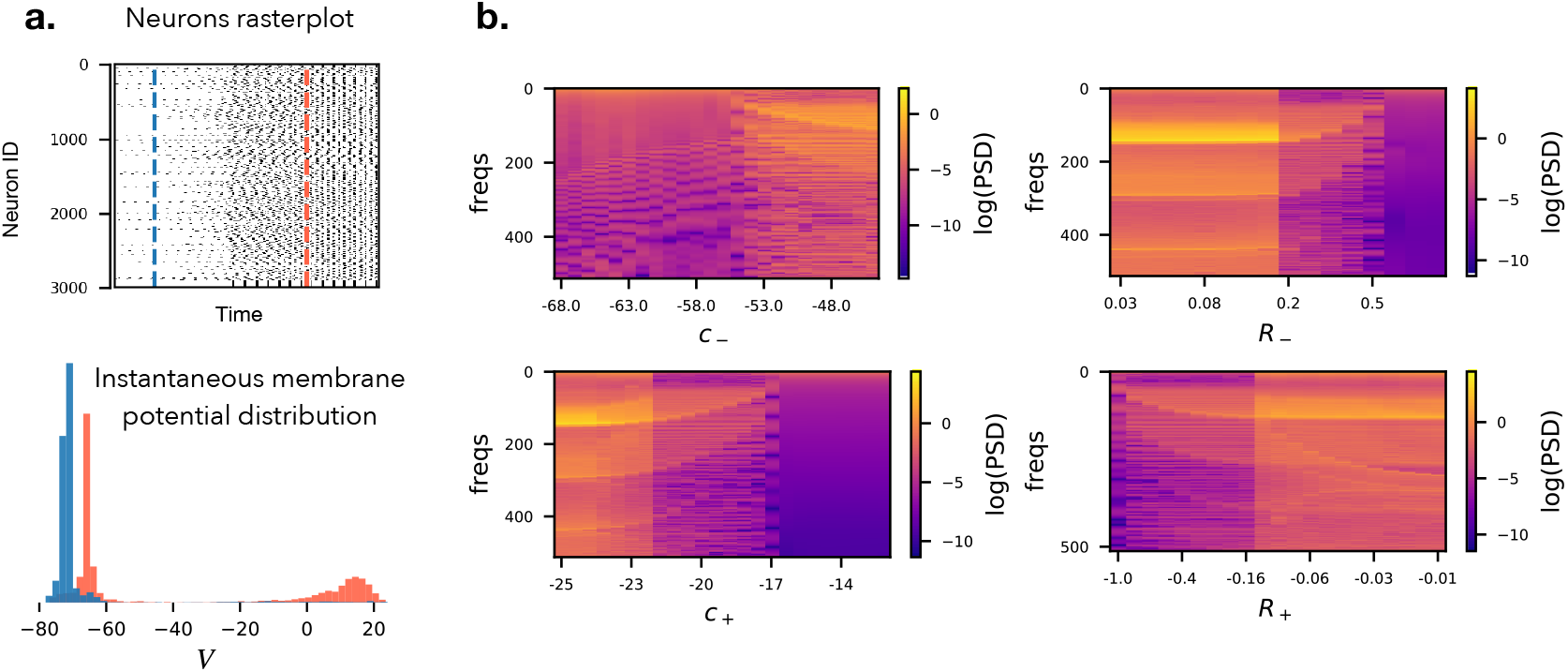
Quadratic approximation of nullcline geometry: **(a)** Neuronal network simulations of all-to-all coupled HH-like neurons can display bimodal distribution of the membrane potential when a subpopulation transitions from sub-to supra-threshold activity. By changing the values of the parabola coefficients (*c*_−_, *R*_−_, *c*_+_, *R*_+_), which describe the nullcline geometry of the model, the results of the mean-field simulation change qualitatively. We show this parameters’ effect by plotting the power spectral density for several values of these parameters (here we used [*K*^+^]_*bath*_ = 15.5). We can observe that for all the parameters, frequency shifts and sudden state transitions occur.

**Fig. S2.**
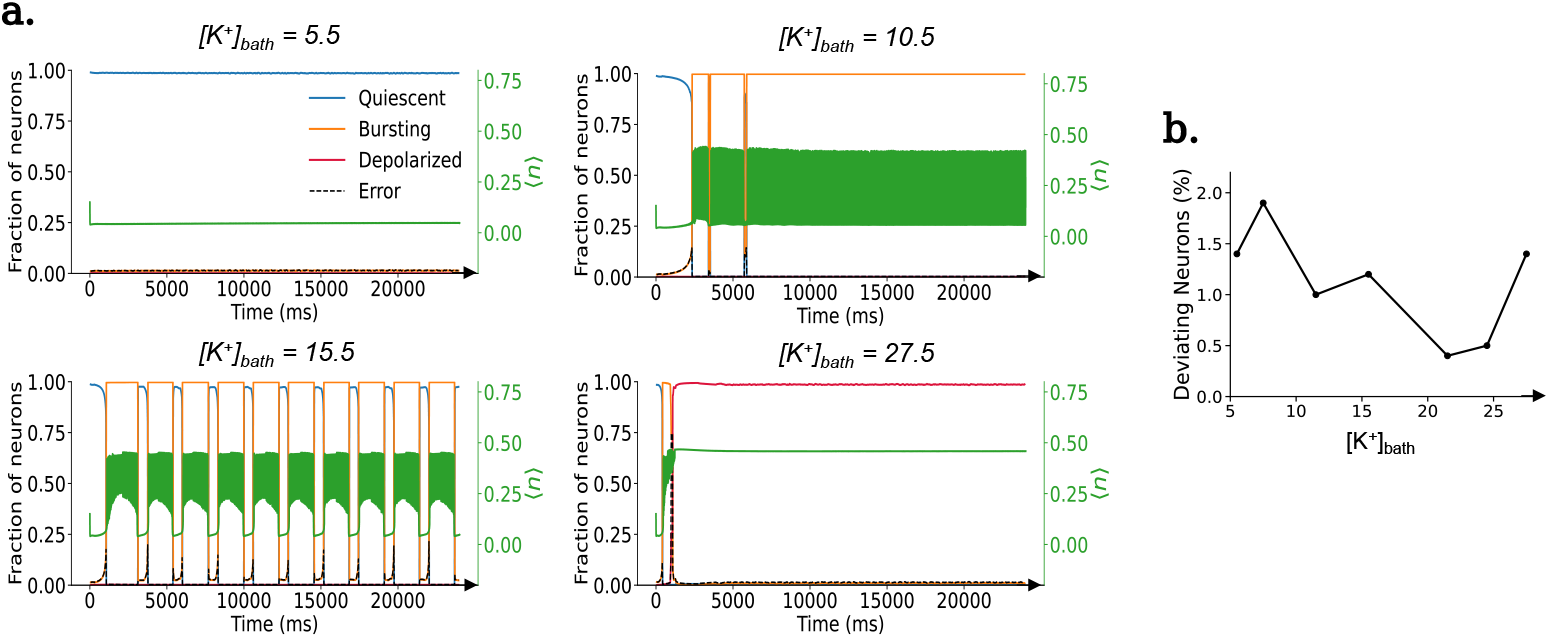
Quantifying the error of the first moment approximation: **(a)** For several [*K*^+^]_*bath*_ values corresponding to different dynamical regimes we simulated the activity of a population of N = 3000 all-to-all coupled HH-type neurons (parameters same as Fig. 3.3 in Table S2) and classified them in three states: Quiscent (Blue), Bursting (Orange) and Depolarized (Crimson red) based on their individual gating variable (*n*) values. Along with that, using the right-hand axis, the population mean of the gating variable is plotted in green. Errors (Black dotted) have been calculated by calculating which fractions of neurons are not following the state of the mean gating variable. **(b)** The error percentage (neurons not following the mean behavior) is plotted against several [*K*^+^]_*bath*_ values.

**Table S2.**
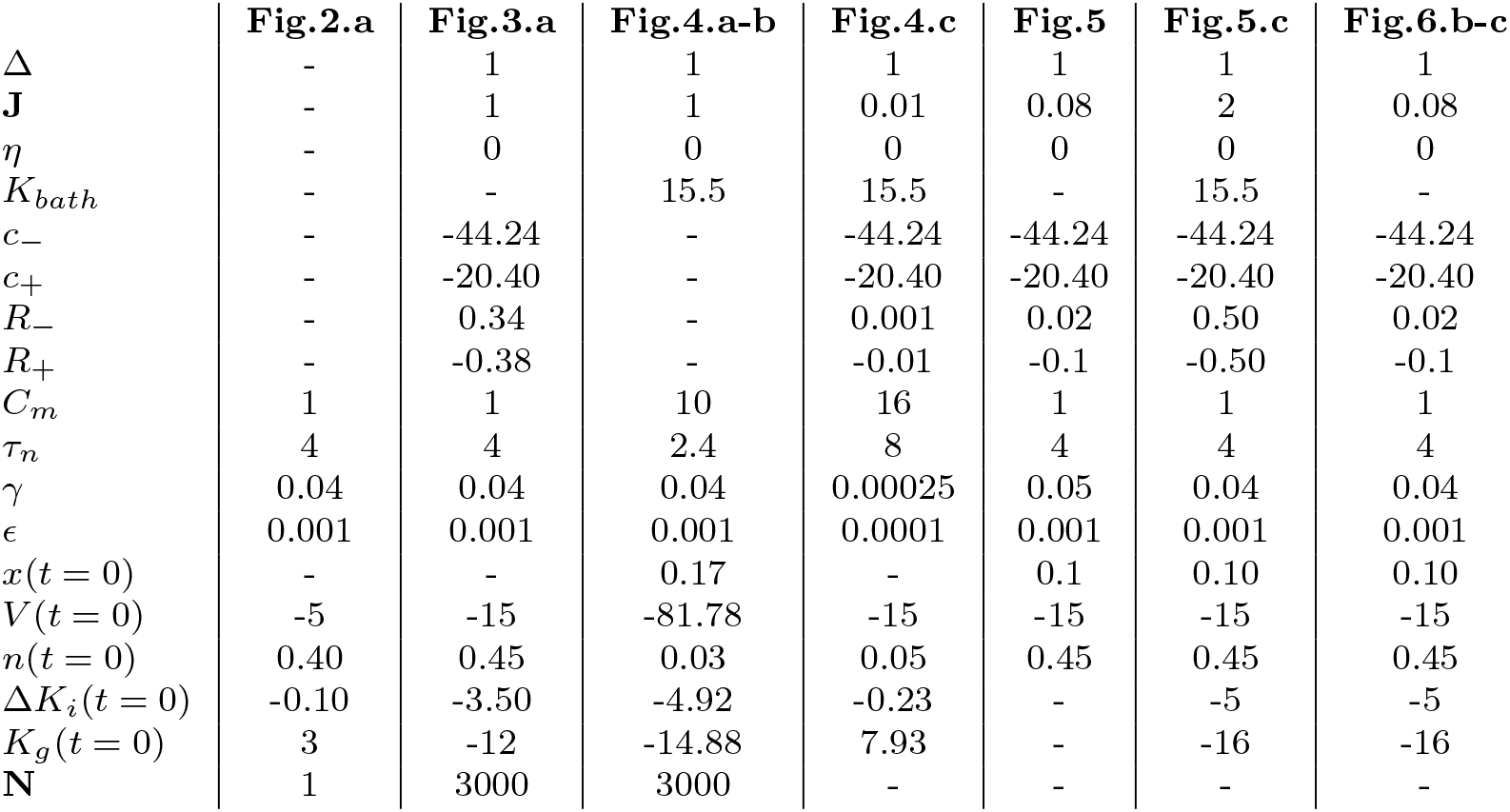
List of parameters’ values used for each simulation in the manuscript. If not present, the parameter is not applicable for that case, or different values have been used, as specified in the corresponding caption.

|  | Fig.2.a | Fig.3.a | Fig.4.a-b | Fig.4.c | Fig.5 | Fig.5.c | Fig.6.b-c |
| --- | --- | --- | --- | --- | --- | --- | --- |
| $\Delta$ | - | 1 | 1 | 1 | 1 | 1 | 1 |
| $J$ | - | 1 | 1 | 0.01 | 0.08 | 2 | 0.08 |
| $\eta$ | - | 0 | 0 | 0 | 0 | 0 | 0 |
| $K_{bath}$ | - | - | 15.5 | 15.5 | - | 15.5 | - |
| $c_-$ | - | -44.24 | - | -44.24 | -44.24 | -44.24 | -44.24 |
| $c_+$ | - | -20.40 | - | -20.40 | -20.40 | -20.40 | -20.40 |
| $R_-$ | - | 0.34 | - | 0.001 | 0.02 | 0.50 | 0.02 |
| $R_+$ | - | -0.38 | - | -0.01 | -0.1 | -0.50 | -0.1 |
| $C_m$ | 1 | 1 | 10 | 16 | 1 | 1 | 1 |
| $\tau_n$ | 4 | 4 | 2.4 | 8 | 4 | 4 | 4 |
| $\gamma$ | 0.04 | 0.04 | 0.04 | 0.00025 | 0.05 | 0.04 | 0.04 |
| $\epsilon$ | 0.001 | 0.001 | 0.001 | 0.0001 | 0.001 | 0.001 | 0.001 |
| $x(t=0)$ | - | - | 0.17 | - | 0.1 | 0.10 | 0.10 |
| $V(t=0)$ | -5 | -15 | -81.78 | -15 | -15 | -15 | -15 |
| $n(t=0)$ | 0.40 | 0.45 | 0.03 | 0.05 | 0.45 | 0.45 | 0.45 |
| $\Delta K_i(t=0)$ | -0.10 | -3.50 | -4.92 | -0.23 | - | -5 | -5 |
| $K_g(t=0)$ | 3 | -12 | -14.88 | 7.93 | - | -16 | -16 |
| $N$ | 1 | 3000 | 3000 | - | - | - | - |

## Notes

### Competing Interest Statement

The authors have declared no competing interest.

### Summary of Updates

This revised version of the manuscript addresses the reviewers comments through targeted changes to both the text and the analysis. We revised several mathematical expressions and accompanying explanations to improve clarity and consistency. The description of the model assumptions and limitations has been expanded and clarified throughout the manuscript. We revised Figure 4. In addition, we replaced the previous Figure 5 with a new figure and corresponding analysis. The new Figure 5 presents an extended bifurcation analysis of the model, providing additional insight into the dynamical regimes and their dependence on model parameters.

